# LSM2-8 and XRN-2 contribute to the silencing of H3K27me3-marked genes through targeted RNA decay

**DOI:** 10.1101/701581

**Authors:** Anna Mattout, Dimos Gaidatzis, Jan Padeken, Christoph Schmid, Florian Aeschlimann, Véronique Kalck, Susan M. Gasser

## Abstract

In fission yeast and plants, RNA-processing pathways contribute to constitutive and facultative heterochromatin silencing, complementing well-characterized pathways of transcriptional repression. However, it was unclear whether this additional level of regulation occurs in metazoans. Here we describe a pathway of silencing in *C. elegans* somatic cells, in which the highly conserved, RNA binding complex LSM2-8 selectively silences heterochromatic reporters and endogenous genes bearing the Polycomb mark H3K27me3. Importantly, the LSM2-8 complex works cooperatively with XRN-2, a 5’-3’ exoribonuclease, and disruption of the pathway leads to mRNA stabilization. This selective LSM2-8-mediated RNA degradation does not target nor depend on H3K9me2/me3, unlike previously described pathways of heterochromatic RNA degradation. Intriguingly, the loss of LSM2-8 coincides with a localized drop in H3K27me3 levels on *lsm-8*-sensitive loci only. Together this defines a mechanism of RNA degradation that selectively targets a subset of H3K27me3-marked genes, revealing an unrecognized layer of regulation for facultative heterochromatin in animals.

## Introduction

The organization of DNA sequences into highly condensed, dark-staining heterochromatin correlates with reduced gene expression (Saksouk et al., 2015; Trojer and Reinberg, 2007; Wenzel et al., 2011). Heterochromatin can be classified as constitutive or facultative heterochromatin. H3K9me3 is the histone modification that characterizes constitutive heterochromatin, which is found most often on non-coding raepetitive elements (Saksouk et al., 2015; Zeller et al., 2016), while Polycomb-mediated trimethylation of H3K27 is the hallmark of facultative heterochromatin. H3K27me3 defines a repressive state that silences genes as a function of temporal and spatial conditions, for instance, during development (Gaydos et al., 2014; Trojer and Reinberg, 2007). Whereas transcriptional repression is believed to be the primary level of regulation through which both constitutive and facultative heterochromatin act, robust pathways that silence at co- and post-transcriptional levels have been documented in fission yeast and plants (Buhler, 2009; Wang et al., 2016).

Over a decade ago it was shown that transcription and noncoding RNAs were involved in the establishment of heterochromatin-mediated repression in fission yeast (Grewal and Elgin, 2007; Moazed, 2009). Counterintuitively, ncRNA transcripts were shown to promote RNAi-mediated assembly of centromeric heterochromatin by providing both small RNAs and a scaffold to recruit chromatin-modifying enzymes through the RITS complex (Buhler, 2009; Noma et al., 2004; Wang et al., 2016). Additionally, RNAi-independent RNA degradation mechanisms that use the exosome were implicated in constitutive heterochromatic gene silencing in *S. pombe* (Buhler et al., 2007). The exosome was also implicated in heterochromatic repeat silencing in *Drosophila* (Eberle et al., 2015), and at centromeric and pericentromeric loci in *Arabidopsis* (Shin et al., 2013). RNA degradation was also suggested to contribute to rDNA stability and subtelomeric silencing in budding yeast (Vasiljeva et al., 2008).

In *S. pombe*, a variety of RNA associated factors were shown to promote H3K9me2/3 silencing in a partially redundant manner, acting through mechanisms that process RNA transcripts. These include HP1(Swi6) (Keller et al., 2012), Red1 and Mmi1 (Egan et al., 2014; Touat-Todeschini et al., 2017; Yamanaka et al., 2013; Zofall et al., 2012), Pla1 (Yamanaka et al., 2013), Pab2 (St-Andre et al., 2010; Yamanaka et al., 2013), and Dhp1/Xrn2 (Chalamcharla et al., 2015; Tucker et al., 2016). These studies document multiple links between RNA factors and constitutive heterochromatin in plants and yeast, yet no compelling parallel has been reported to date for facultative heterochromatin in animals. Using a genome-wide derepression screen, we have uncovered and characterized such a pathway in embryos and differentiated tissues of *C. elegans*.

In an earlier genome-wide RNAi screen that monitored the derepression of an integrated heterochromatic reporter in *C. elegans*, we identified 29 factors that were essential for heterochromatin silencing in embryos (Towbin et al., 2012; Fig. 1A,B). While most of the validated hits were chromatin modifiers and transcription-related proteins (Towbin et al., 2012), among them were three subunits of the RNA-binding Like-SM (LSM) complexes (*gut-2/lsm-2, lsm-5* and *lsm-6*). LSM proteins are highly conserved throughout evolution, with the *C. elegans* proteins sharing up to 94% homology with their human counterparts (Fig. S1A). The two LSM complexes, LSM1-7 (cytoplasmic) and LSM2-8 (nuclear), were shown to function in RNA metabolism and splicing, but not in transcription *per se* (Beggs, 2005; Cornes et al., 2015; Golisz et al., 2013; Kufel et al., 2004; Perea-Resa et al., 2012; Tharun, 2009). The cytoplasmic LSM1-7 complex partners with decapping enzymes, which render RNA sensitive to the 5’ to 3’ XRN-1 exonuclease activity, while the LSM2-8 complex has been proposed to work with XRN-2 to promote nuclear RNA decay (Beggs, 2005; Tharun, 2009). No role for LSM proteins in heterochromatin silencing has been previously reported.

**Figure 1:**
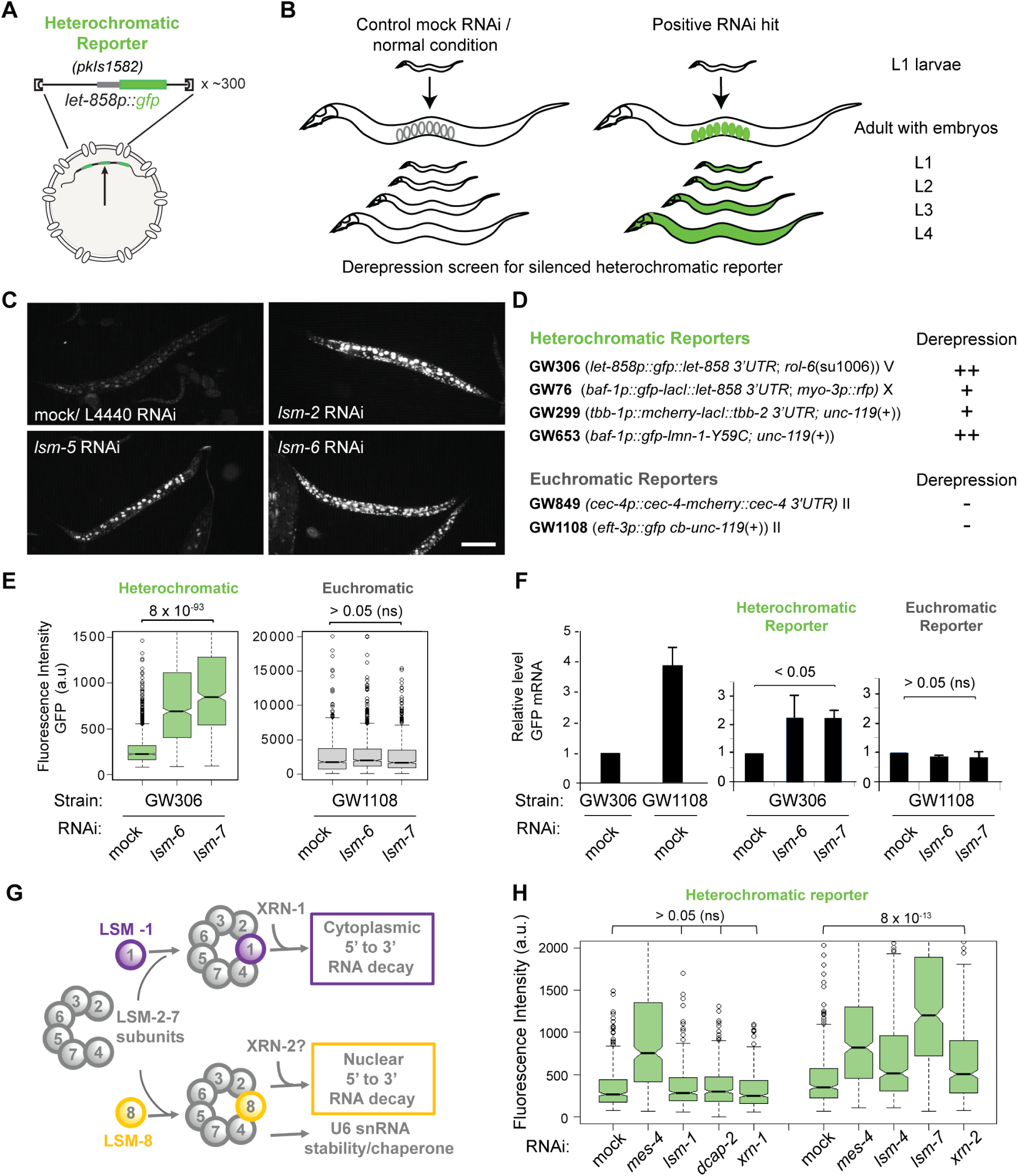
LSM proteins silence heterochromatic reporters, but not euchromatic reporters. **A,** Sketch of the integrated, high-copy number heterochromatic reporter *pkIs1582* from strain GW306 used in the genome-wide screen (Towbin et al., 2012). **B,** Here RNAi-based derepression was monitored in progeny of all stages by increased GFP fluorescence. **C,** Fluorescence microscopy of *pkIs1582*-encoded GFP in L4 larvae with indicated RNAi versus control (mock/L4440). Bar, 100 μm. **D,** Heterochromatic and euchromatic reporters scored by eye for derepression (+, ++: strong and very strong derepression, respectively) upon LSM RNAi (Table S1). **E,** Quantitation of derepression in L1 larvae by the worm sorter following indicated RNAi. Box plots of fluorescence intensity in arbitrary units (a.u), with whiskers = 1st and 3rd quartiles, black lines: median, black circles outliers. n =2000, 1068, 613 and 875, 111, 1026. Indicated p-values by Student’s t test, (n.s.= p> 0.05). **F,** qPCR analysis of GFP mRNA in L1 larvae as in (E), normalized to *his-56* and *pmp-3* mRNA. GFP from GW306 strain is set as 1 (left), and mock RNAi conditions are set as 1 (right). N= 2, n=3, bars (mean ± s.d). **G,** The two main LSM complexes and functions (Beggs, 2005; Tharun, 2009). **H,** GFP fluorescence of the heterochromatic reporter (*pkIs1582;* GW306) in L1 larvae after RNAi treatment for indicated genes. n= 396 for each treatment.

Our initial screen implicated LSM-2, LSM-5 and LSM-6 in gene silencing based on robust GFP derepression of a heterochromatic reporter in worm embryos, after RNAi-mediated knockdown of the individual genes (Fig. 1A,B). In this de-silencing assay, the heterochromatic reporters consisted of an integrated array of several hundred copies of a plasmid, carrying a GFP-encoding reporter gene driven by a ubiquitously expressed promoter that was repressed in wild-type *C. elegans*. In a copy-number dependent manner, these heterochromatic reporters acquire the histone modifications H3K9me2/3 and H3K27me3 and are sequestered at the nuclear envelope, thereby mimicking endogenous heterochromatin (Meister et al., 2010; Towbin et al., 2010; Wenzel et al., 2011). Loss of either of the two H3K9 methyltransferases (MET-2, SET-25) or the EZH2 homolog, MES-2, which methylates H3K27, led to reporter derepression (Towbin et al., 2012).

Here we examined in depth the pathway by which LSM proteins contribute to heterochromatic silencing in *C. elegans*. We find that the LSM2-8 complex works with XRN-2 in a post-transcriptional RNA decay pathway that selectively targets transcripts arising from endogenous genes enriched for H3K27me3. The LSM1-7 complex is not involved in this silencing pathway. LSM8-mediated silencing can occur at all developmental stages, and in all somatic cells. Not only are the LSM8*-*sensitive loci preferentially enriched for Polycomb-mediated H3K27me3, but the level of H3K27me3 on these genes drops in animals lacking *lsm-8*. This argues for a feedback loop, in which LSM2-8 serves as an intermediary that triggers the degradation of specific transcripts while concomitantly enhancing a repressive chromatin state. Upon the loss of *lsm-8,* the animals are sterile, have premature death phenotypes and misexpress the HOX gene *egl-5*. This is the first molecular pathway of RNA degradation in any organism that is correlated with the characteristic histone modification of facultative heterochromatin, H3K27me3. It shows that the selective degradation of heterochromatic transcripts in the nucleus can supplement repression on the level of transcription to silence target genes in animals.

## Results

### LSM proteins selectively silence heterochromatic reporters throughout somatic differentiation

The initial genome-wide RNAi screen for factors involved in heterochromatin silencing monitored *let-858p::gfp* derepression from the heterochromatic reporter *pkIs1582* (strain GW306), in embryos, (Fig. 1A,B; Table S1; Towbin et al., 2012). We used RNAi against *lsm-2*, *lsm-5* and *lsm-6* to analyze the derepression/ increased expression of GFP throughout *C. elegans* development. We found a reproducible and robust derepression at all stages *i.e.* in embryos, L1 to L4 larval stages and in adult worms (Fig. 1C, Fig. S1B). Moreover, as shown by fluorescence microscopy, elevated levels of GFP were found in nearly every somatic cell type (Fig. 1C, Fig. S1B).

The ubiquitous derepression of the heterochromatic reporter allowed us to quantify GFP expression following RNAi using the COPAS Biosorter. The Biosorter uses flow cytometry to quantify fluorescence of single worms with high throughput, generating robust population-wide measurements. We find highly significant upregulation of GFP in populations of L1 larvae following RNAi for *lsm-2, lsm-5* and *lsm-6,* as compared to the control/mock RNAi treated worms (L4440; Fig. S1C). The differences in derepression between *lsm-2, lsm-5* and *lsm-6* most likely reflect the different knock-down efficiencies of the respective RNAi clones (Fig. S1C). The population-wide effect of *lsm-2* RNAi was as strong as RNAi against the HMT MES-4, a positive control that ablates H3K36 methylation (Towbin et al., 2012).

Given the various RNA processing functions attributed to LSM proteins (Tharun, 2009), we asked whether the derepression we scored upon *lsm* depletion acted uniquely on the heterochromatic reporter used. We tested *lsm* RNAi on the expression of four independent heterochromatic reporters, each with a distinct combination of promoter, reporter gene (encoding GFP or mCherry), 3’ UTR, and site of integration (Fig. 1D, Table S1). We also tested for effects of *lsm-2, lsm-5* or *lsm-6* RNAi on two euchromatic reporters respectively from the strains GW849 and GW1108, which carry single copy transgenes (encoding GFP or mCherry) integrated into non-heterochromatic regions of the genome. Whereas all the heterochromatic reporters tested showed significant derepression (Fig. 1D,E, Fig. S1C), the euchromatic reporters showed no change in expression following *lsm* RNAi (Fig. 1D,E, Fig. S1F-G). This specificity was confirmed not only for RNAi against *lsm-2, -5* and *-6,* but also for *lsm-7* RNAi (Fig. S1B,D), another subunit of the LSM complexes.

The lack of response observed for the euchromatic reporters does not reflect differences in the basal or background fluorescence level, given that the four heterochromatic reporters, despite having different initial GFP expression levels, were all derepressed following *lsm-7* RNAi (Fig. S1D). Moreover, neither *lsm-6* nor *lsm-7* RNAi had an effect on a highly expressed single copy transgene (*eft-3p::gfp*, GW1108), whose GFP signal is higher than the repressed, *let-858p::gfp* heterochromatic reporter (from the strain GW306; Fig. 1E), nor on the euchromatic *cec-4p::mCherry* reporter in GW849, which is expressed at a lower level than this heterochromatic reporter (Fig. S1F,G).

Finally, to make sure that the increased expression reflects changes in mRNA and not altered protein synthesis or turnover, we monitored the relative increase in *gfp* mRNA levels in GW306 and GW1108 by qPCR. Indeed, the heterochromatic *let-858p::gfp* reporter showed higher steady-state levels of mRNA following *lsm-6* and *lsm-7* RNAi, while the euchromatic *eft-3p::gfp* mRNA did not (Fig. 1F). Taken together, this suggests that the *C. elegans* LSM factors regulate mRNA levels, rather than protein turnover or translation, to silence specifically reporters with heterochromatic features. This occurs both during somatic cell differentiation and in post-mitotic cells throughout development.

### RNAi implicates LSM2-7 and XRN-2, but not LSM-1 and XRN-1, in reporter repression

The LSM proteins 2 through 7 are shared subunits of two related complexes: the LSM1-7 complex is primarily cytoplasmic, while the LSM2-8 complex is nuclear (Beggs, 2005; Fig. 1G). The two complexes also differ in their co-factors. The cytoplasmic LSM1-7 complex acts together with the 5’→3’exoribonucleases, XRN-1 and the decapping enzymes DCAP-1 and DCAP-2 to mediate cytoplasmic RNA decay (Tharun, 2009). The LSM2-8 complex on the other hand was suggested to work in concert with the nuclear 5’→3’exoribonucleases XRN-2 (Tharun, 2009). To determine which of the two LSM complexes contribute to heterochromatic gene silencing, we compared reporter derepression after RNAi against *lsm-1*, which is unique to the LSM1-7 complex, with the levels after RNAi against two shared subunits, *lsm-4* and *lsm-7.* Strong heterochromatic reporter derepression was scored upon knockdown of *lsm-4* and *lsm-7*, while no effect was observed following RNAi against *lsm-1* (Fig. 1H). RNAi efficiency was similar in *lsm-1* and *lsm-7* RNAi treated larvae (Fig. S1E). In addition, RNAi against the LSM1-7 associated factors, *dcap-2* and *xrn-1*, failed to provoke heterochromatic reporter derepression, while RNAi against *xrn-2* did (Fig. 1H). In summary, our RNAi studies argued that knockdown of the related LSM proteins, LSM-2, -4, -5, -6, and -7, but not of LSM-1 or its associated factors, triggered heterochromatin reporter derepression.

### Deletion of *lsm-8* leads to efficient derepression, while *lsm-1* or *dacp-2* deletions do not

The failure of *lsm-1* RNAi to derepress the reporter suggested that the LSM1-7 complex is not involved in heterochromatic silencing, and suggested that the LSM2-8 complex is. Because attempts to use RNAi to downregulate *lsm-8*, the only unique subunit of LSM2-8, were ineffective in our hands, we generated a full *lsm-8* deletion by CRISPR/Cas9, replacing the *lsm-8* locus with a red fluorescent marker gene with pharynx-specific expression. This allows tracking of the null allele (Fig. 2A). We found that somatic development the homozygous *lsm-8* ^-/-^ worms - including gonad formation - was similar to wild-type worms up through the L3 and L4 larval stages (Fig. S2A,B), yet the adult homozygous mutant was sterile. At the young adult stage (after L4) the gonads in *lsm-8 ^-/-^* animals became abnormal and failed to support oocyte maturation (Fig. S2C). To stably maintain the *lsm-8* deletion we balanced the deletion with the *nT1*[*qIs51*] balancer, which expresses a GFP marker in the pharynx. This allowed us to sort *lsm-8^-/-^* homozygous from heterozygous *lsm-8 ^+/-^* worms for further analysis because heterozygous *lsm-8*^-/+^ worms would have both red and green fluorescence in the pharynx, and homozygous *lsm8*^-/-^ worms express only red (Fig. 2B).

**Figure 2:**
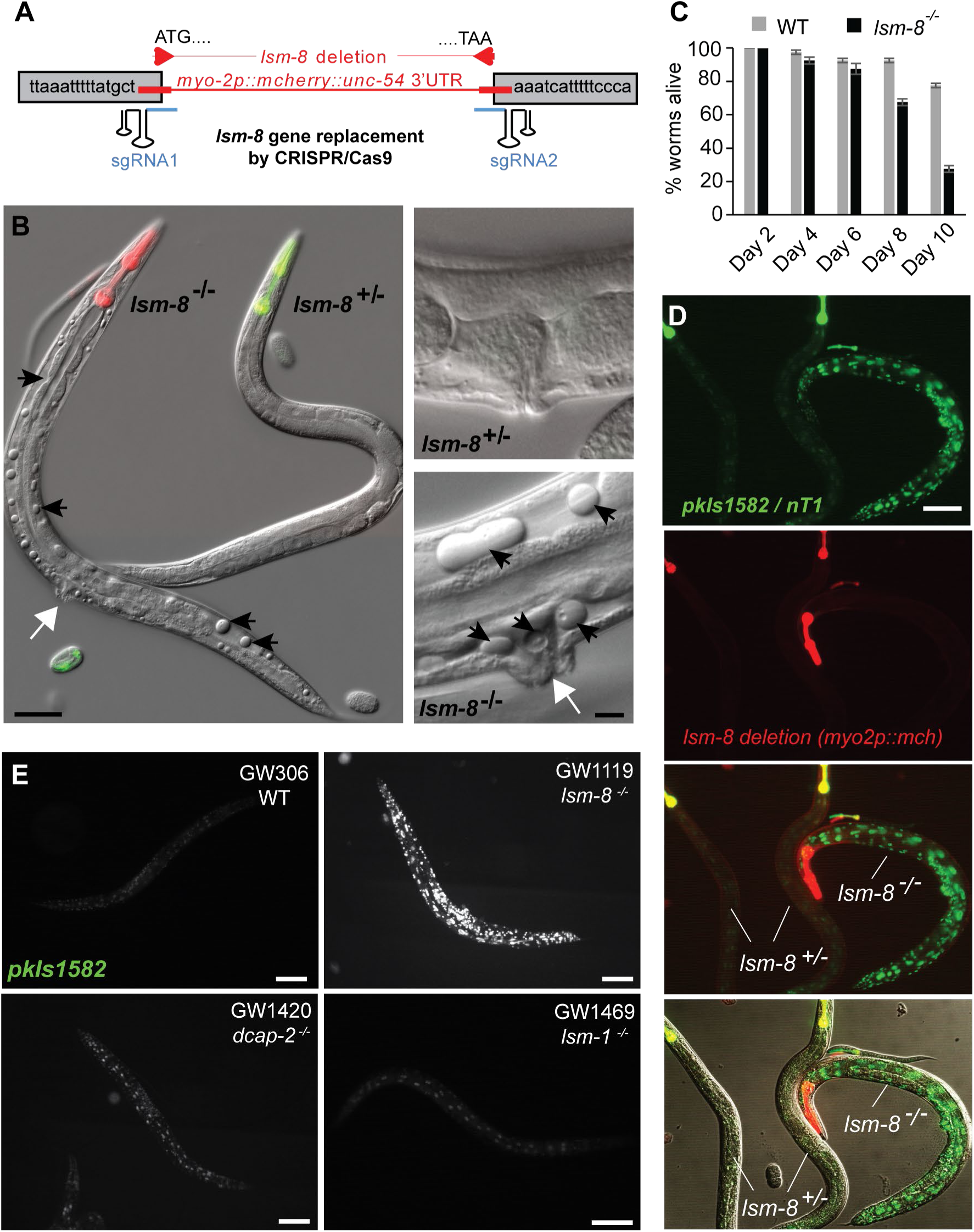
LSM2-8 mediates heterochromatic silencing, and prevents sterility and premature death. **A,** Schematic view of the *lsm-8* deletion/gene replacement created by CRISPR-Cas9. **B,** Differential interference contrast (DIC) images of young adults (GW1120) merged with pharynx fluorescence to identify genotypes, as in (D). *lsm-8*^-/-^ worms accumulate cavities and vacuoles (black arrows), and protruding vulva (white arrows). Right, enlargement of the vulva region. Bars, 50 μm (left) and 10 μm (right). **C,** Survival assay at 22.5°C after hatching shows premature death of *lsm-8^-/-^* worms. N=4, n = 40 per genotype. Bars (mean ± s.d). **D,** View of *lsm-8*^+/-^ (yellow pharynx in merge) and *lsm-8*^-/-^ (red pharynx only) worms carrying the *pkIs1582* heterochromatic reporter. Red and green channels are shown separated and merged. Bar, 100 μm. **E,** Heterochromatic reporter *pkIs1582* derepression in *lsm-8^-/-^* background compared to the WT, *dcap-2^-/-^* and *lsm-1^-/-^* background level. Bars, 100 μm.

Although the *lsm-8*^-/-^ animals developed to adulthood, they had protruding vulva, showed empty cavities or vacuoles in differentiated tissues, died prematurely and were 100% sterile (no oocytes; Fig. 2B, Fig. S2C). The presence of cavities coincides mainly with premature death in adult stages (Fig. 2B). Unlike *lsm-1* mutants (Cornes et al., 2015), worms lacking *lsm-2* or *lsm-5,* are phenotypically similar to *lsm-8*^-/-^ mutants, with protruding vulva, abundant vacuoles and 100% sterility (Fig. S2C).

Using pharyngeal fluorescence to distinguish homozygous *lsm8*^-/-^ from heterozygous *lsm8*^+/-^ animals in the F1 progeny, we monitored the repression of the heterochromatic reporter *pkIs1582* integrated in the strain carrying the *lsm-8* null allele. Derepression was as strong as with *lsm-7* RNAi and was restricted to homozygous *lsm-8^-/-^* animals (red pharynx; Fig. 2D). To be certain that the phenotype was specific for the LSM2-8 complex, we obtained and backcrossed homozygous *lsm-1^-/-^* and *dcap-2^-/-^* animals, coupling these genomic deletions with the same heterochromatic reporter. While the *lsm-8^-/-^* larvae had uniformly GFP-positive nuclei due to reporter derepression throughout the animal, none of the *lsm-1* and *dcap-2* null animals had GFP signals above the wild-type (WT) background levels (Fig. 2E). We conclude that the loss of heterochromatic silencing stems from loss of a functional LSM2-8 complex, while there is no indication that the LSM1-7/DCAP-2 pathway of cytoplasmic RNA degradation by XRN-1 is involved.

Given the sterility and severe germline phenotype that was manifest in the *lsm-8^-/-^* animals, we next examined derepression of the reporter within the gonad. Whereas the heterochromatic reporter *pkIs1582* was derepressed in the somatic gonadal cells (distal tip cell, gonadal sheath, and spermathecal cells), as well as in nearly all somatic cells in L4 larvae both after *lsm-7 RNAi* and in *lsm-8^-/-^* worms, the germline itself (germ cells, in the dashed red line) had no GFP expression (Fig. S2D-F). We wondered if this might reflect redundancy with the piRNA pathway, which mediates germline specific silencing (Shirayama et al., 2012). However, there was again no derepression in the *lsm8*^-/-^ germ cells (GW1119) when the mutation was coupled with RNAi against the piRNA-related factor, *csr-1* (Fig. S2G), or *prg-1* (data not shown). Thus LSM2-8 pathway seems to affect primarily reporter expression in somatic cells.

### LSM2-8 is required to maintain silent endogenous heterochromatin

To examine changes in endogenous transcript levels provoked by loss of LSM-8, we performed a strand-specific total RNA-seq on WT and homozygous *lsm-8^-/-^* sorted L3 larvae (Fig. 3A). In order to compare the LSM2-8 silencing pathway with the well-characterized pattern of repression mediated by H3K9 methylation (Towbin et al., 2012; Zeller et al., 2016), transcriptome data from sorted *lsm-8^-/-^* L3 larvae was compared with data obtained in parallel from larvae carrying the double deletion *met-2^-/-^ set-25^-/^*^-^ (which lack all detectable H3K9me), or the triple mutation, *met-2^-/-^ set-25^-/-^; lsm-8^-/^*.. In each case, the WT, single, double and triple mutant worms were sorted by fluorescence (no pharyngeal green signal indicating *lsm8^-/-^*) and by size, to generate uniform populations of L3 stage larvae for each genotype (Fig. 3A). Slight shifts in timing between samples can occur even among larvae at the same developmental stage due to the time required for the sorting process. Therefore, we chose and matched pairwise the samples from the four genotypes that were the closest in timing. We scored for developmental timing using a characteristic temporal fluctuation of a subset of somatic genes that were shown to be robust markers for developmental synchrony (Hendriks et al., 2014) (Fig. S3A).

**Figure 3:**
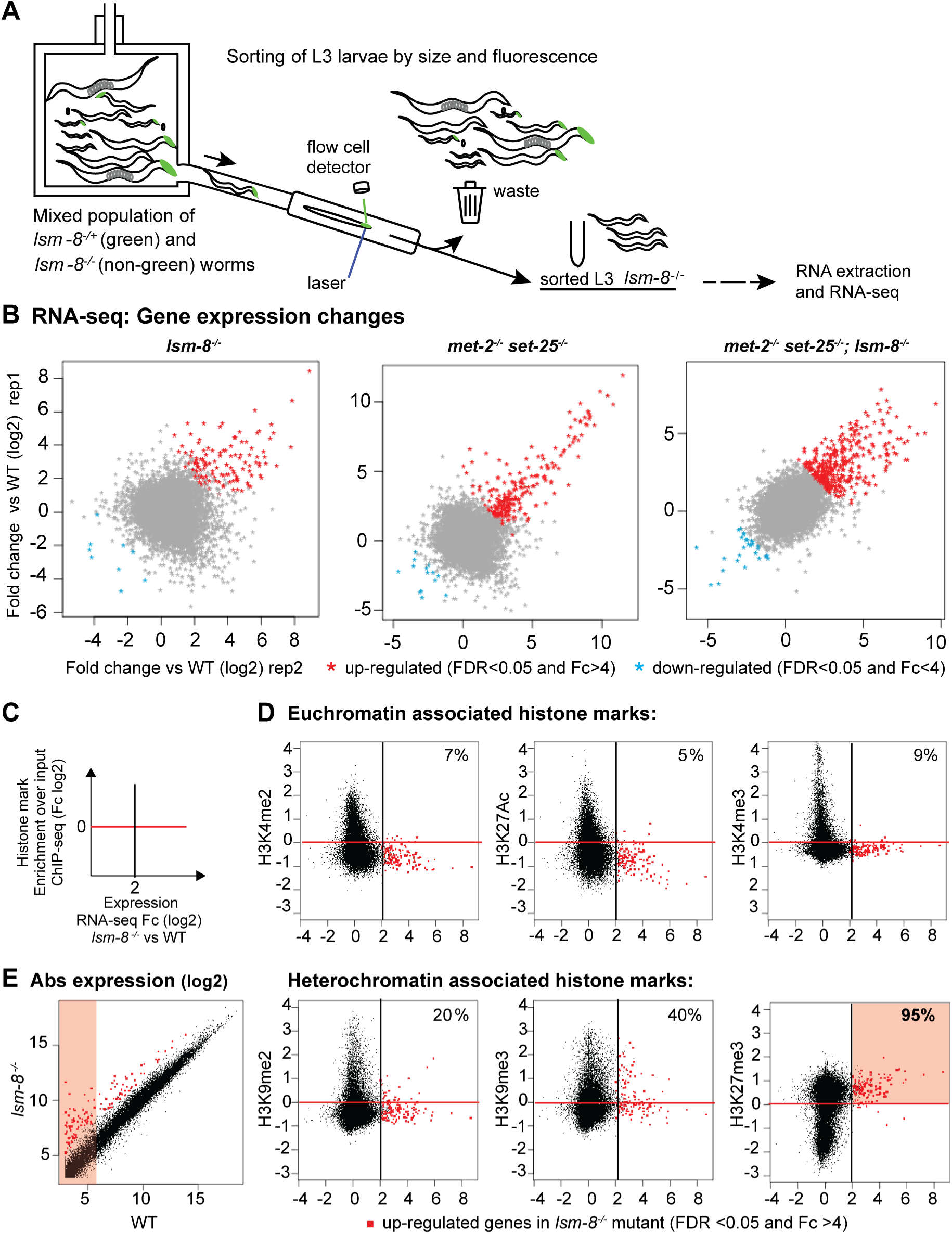
95% of genes silenced by LSM2-8 carry the Polycomb mark H3K27me3. **A,** Worm sorting process. L3 worms with the four following genotypes: *lsm-8^-/-^*; *met-2^-/-^ set-25^-/-^*; triple mutant and WT were sorted and harvested using the same criteria. **B,** Deletion of *lsm-8* (*lsm-8^-/-^*) derepresses significantly >100 genes (FDR <0.05 and Fc >4). Relative gene expression profiles are shown as scatter plots, with Fold-change (Fc) in log2 (log2) for two RNA-seq replicas of L3 sorted worms of the indicated genotype versus WT. Each dot corresponds to a gene. **C,** Scatter plot that compares the average gene expression changes in *lsm-8^-/-^* worms (x-axis in log2, RNA-seq L3 stage) versus enrichment for a histone modification (y-axis in log2, ModEncode data of WT L3 stage). Up-regulated genes (FDR >0.05 and Fc >4) in the *lsm-8^-/-^* mutant are in red to the right of the black line, and genes enriched for the histone mark are above the red line (enriched over input). **D,** Scatter plots as (C), with each dot representing a gene. Upper row, euchromatic marks; lower row, heterochromatin marks. % indicates genes in upper right zone: LSM-8 regulated and enriched genes for indicated mark. **E,** Scatter plot of absolute gene expression (normalized reads count, log2) of *lsm-8^-/-^* versus WT. Red dots as in (D). Values under 6 (log2) are considered to have very low expression (pink shading).

Deletion of *lsm-8* led to the up-regulation of transcripts of 122 genes (false discovery rate (FDR) <0.05 and fold change (Fc) >4), while only 9 genes were down-regulated (Fig. 3B, Table S2). Using less stringent cut-off values (Fc >2), there were 1332 genes selectively upregulated in *lsm-8^-/-^* larvae. We confirmed that expression differences found between the mutant and WT worms (FDR <0.05 and Fc >4) cannot be attributed to the slight differences in timing between samples, since the expression changes that stem from differences in developmental timing (rising genes, Fig. S3B) did not reflect the genes upregulated in *lsm-8 ^-/-^* L3 larvae or in the other mutants (Fig. 3B).

We looked for additivity or epistasis between the LSM2-8 and H3K9me repression pathways by comparing genome-wide RNA-seq data from the single, double and triple mutants. While the loss of H3K9 methylation (*met-2 set-25)* led to the strong upregulation of 219 genes (FDR <0.05 and Fc >4), only 36% of these overlapped with those derepressed in the *lsm-8^-/-^* mutant (Fig. S3C, yellow; Table S2). In addition, there are large subsets of derepressed genes that are upregulated exclusively in the *lsm-8* or the *met-2 set-25* mutant, arguing that the pathways are largely independent (Fig. S3C, blue and pink shading). Consistent with this hypothesis, the triple mutant (*lsm 8^-/-^; met-2^-/-^ set-25^-/-^*) showed little epistasis: an even larger number of genes were strongly derepressed (FDR <0.05 and Fc >4) in the triple mutant than in either *lsm-8* or the *met-2 set-25* strains (367 genes vs 122 and 219 respectively; Table S2, Fig. S3D). To illustrate this additivity, we selected a group of genes that were mildly upregulated upon loss of either *lsm-8* or *met-2 set-25* (Fc <2; red boxed area in graph Fig. S3D) and examined their behavior in the triple mutant. The vast majority were derepressed >4-fold in the triple mutant (orange spots in Fig S3D, Table S2). Consistent with this, *gfp* expression from the heterochromatic reporter, which bears both H3K9 and H3K27 methylation, also showed an additive relationship (Fig. S3E). We conclude that the LSM2-8 pathway of silencing for endogenous loci is distinct from that mediated by H3K9 methylation and does not depend on H3K9me3, even though some genes were targeted by both pathways.

The additivity of phenotypes extends beyond gene expression. As indirect support for LSM2-8 and H3K9me acting on parallel pathways, we note that unlike the *lsm-8* deletion which strongly enhanced premature lethality during somatic growth over 10 days, the loss of H3K9 methylation alone does not (Fig. S3F), but rather slows development in a stochastic manner (Zeller et al., 2016). As expected, in worms bearing the combination of *met-2 set-25* with the *lsm-8* null allele, we observed enhanced lethality, i.e., the mutations were not epistatic, but additive (Fig. S3F).

### Over 93% of LSM2-8 silenced genes bear H3K27 trimethylation

Given that heterochromatic reporters, but not euchromatic reporters, are upregulated upon loss of *lsm-8,* we examined whether the genes upregulated by loss of LSM-8 share a common set of histone modifications. To do so, we plotted our L3 RNA-seq data against the normalized ChIP-seq data for common histone modifications found the genome of WT L3 larvae, generated by ModEncode (Table S3, Fig. 3C-E, Fig. S4A). In worms, as in most organisms, H3K4me2, H3K4me3, and H3K27ac are associated with active genes (Ho et al., 2014; Liu et al., 2011; Wenzel et al., 2011), while H3K9me2/3 and H3K27me3 are repressive histone marks that co-localize with heterochromatin (Ahringer and Gasser, 2018; Wenzel et al., 2011). We found that the genes strongly derepressed in the *lsm-8^-/-^* mutant (Fc>4 and FDR<0.05), were depleted for the active marks, as well as for H3K9me1 in WT larvae (Fig. 3C-D; Fig. S4A). When *lsm-8-*upregulated genes were plotted against their relative enrichment for repressive marks, we found a striking correlation of LSM-8 sensitivity with the repressive Polycomb mark, H3K27me3. Over 95% of the genes that were derepressed in the *lsm-8^-/-^* mutant were enriched for H3K27me3 (Fig. 3D, Fig. S4A). This was true not only for the genes that met our most stringent cut-off values (Fc>4 and FDR <0.05), but for the genes upregulated between 2- and 4-fold by *lsm-8* ablation (Fig. 3D, Tables S2-S5). In contrast, only 20% of the targeted genes were enriched for H3K9me2, a value that matches the genome-wide presence of H3K9me2-marked genes in L3 larvae, and roughly 40% of the upregulated genes carried H3K9me3 (Fig. 3D).

H3K27me3 and H3K9me3 ChIP-seq signals co-localize more frequently over the *C. elegans* genome, than they do in mammals or flies (Ho et al., 2014). We therefore examined the relationship of H3K27 and H3K9 methylation by examining to which extent the marks colocalize on LSM-8-silenced genes. First, we note that in WT L3 larvae, 41% of H3K27me3-marked genes genome-wide, also bear H3K9me3 (ModEncode). We find a similar rate of H3K9me3 on LSM2-8 targets (40-43%), among which 100% are also H3K27me3 positive (Fig. 3D, Fig. S4A; Tables S2-S4). Thus, the presence of H3K9me3 on LSM-8 target genes simply reflects the rate at which the two marks coincide genome-wide, while the enrichment of H3K27me3 on *lsm-8* sensitive genes in L3 larvae (95%) is highly significant (p<4.2e^-^24). Taken together, we conclude that the histone mark that characterizes LSM2-8-regulated genes is H3K27me3. Consistent with its role in Polycomb-mediated repression (Conway et al., 2015; Grossniklaus and Paro, 2014; Liu et al., 2011; Margueron and Reinberg, 2011), we found that most of the genes significantly upregulated by the loss of LSM-8, are genes that have very low steady-state expression levels in WT worms (Fig. 3E, Fig. S4B).

In further analysis we asked whether the *lsm-8*-sensitive genes had any specific distribution along *C. elegans* chromosomes. This is particularly relevant, given that constitutively repressed H3K9me2/me3 heterochromatin is enriched on chromosomal arms and depleted from chromosome cores (Ahringer and Gasser, 2018). In contrast to this, much like the distribution of the H3K27me3 mark itself (Ho et al., 2014), the genes sensitive to the loss of *lsm-8* were found both in autosomal core regions and along chromosome arms (Fig. S4C). We asked whether other pathways of repression at the L3 larval stage show a similar preference for H3K27me3-marked genes. Performing a similar analysis of chromatin marks on cell cycle genes regulated by the Rb-like repressor, LIN-35, and on developmental genes repressed by PRG-1, a PIWI protein, showed no bias toward H3K27me3 modification (*prg-1* pathway) or even depletion for this mark for the *lin-35* pathway (Fig. S4D). This reinforces our argument that the link of LSM2-8 to Polycomb-marked genes is specific. Interestingly, GO term analysis suggests that the genes silenced by LSM-8 are enriched for genes expressed during the innate immune response, body morphogenesis and cell shape regulation (Table S5), processes that were found to be regulated by Polycomb in other species.

To assess whether LSM2-8 targets H3K27me3-marked genes in other developmental stages than L3 larvae, we performed total RNA-seq on synchronized and sorted WT and homozygous *lsm-8^-/-^* at the L1 larval stage. Deletion of *lsm-8* led to the up-regulation of transcripts of 151 genes (FDR <0.05 and Fc >4), while 59 genes were down-regulated (Fig. S4E, Tables S2, S4). Using Fc >2, 1501 genes were upregulated in *lsm-8^-/-^*. We find a significant (p< 1.17e^-12^) but small subsets of genes (22) that are upregulated upon LSM-8 ablation in both L1 and L3 larval stages and those genes seem to be involved in the immune response (Fig. S4F, Tables S2, S4). Importantly, the L1 stage genes regulated by LSM2-8 are significantly depleted for histone marks associated with euchromatin and are significantly (p< 2.2e^-25^) enriched for the Polycomb mark, H3K27me3 (Fig. S4G, Table S3). Notably, 93% of the genes silenced by LSM2-8 in L1 larvae are found to carry the Polycomb mark.

### Cell-specific HOX gene up-regulation by *lsm-8* ablation in specific cell types

Across all multicellular species, the methylation histone H3K27 by the PRC2 complex leads to cell-type or stage-specific repression of genes implicated in development (Conway et al., 2015; Grossniklaus and Paro, 2014; Liu et al., 2011; Margueron and Reinberg, 2011; Patel et al., 2012), In *C. elegans*, PRC2 consists of MES-2/E(z)/EZH2, MES-3, and MES-6/Esc (Gaydos et al., 2014; Ketel et al., 2005; Yuzyuk et al., 2009). We asked whether the loss of LSM-8 reduced the expression of the Polycomb homologs, impairing catalysis of H3K27me3. However, RNA-seq data in *lsm-8* deficient worms showed no drop in *mes-2, mes-3* or *mes-6* expression, as well as no drop in the expression of *sor-1* and *sop-2* the PRC1-like factors (Table S2), ruling out this potential indirect mechanism.

Intriguingly, HOX genes, the canonical targets of H3K27me3, were not among the strongly upregulated genes in the *lsm-8^-/-^* transcriptome. We reasoned that the transcripts of such genes might not be detected by RNA-seq of *lsm-8* larvae, if their expression occurred only in a limited number of cells. Indeed, cell-type specific repression / derepression events are easily masked in RNA-seq datasets of whole animals. In *C. elegans*, the best conserved HOX cluster repressed by Polycomb includes *lin-39* (essential for development of the vulva), *ceh-13*, *mab-5* and *egl-5* (HOX5/Scr; HOX1/Lab; HOX6-8/Antp; HOX9-13/Abd-B families, respectively) (Hench et al., 2015). Consistently, worms deficient for MES-2, MES-3 or MES-6 (the PRC2 complex) exhibit ectopic HOX gene expression, albeit in a limited number of cells (Ross and Zarkower, 2003). We chose to examine more closely the HOX gene *egl-5,* because even though it did not make our stringent 4-fold cut-off for derepression, it was derepressed to a low level in all *lsm-8* replicates (log2 Fc = 0.32).

The EGL-5 protein is expressed in the tail regions of both hermaphrodites and males, and is required for aspects of male sexual differentiation, especially in male tail development (Ferreira et al., 1999; Ross and Zarkower, 2003). We used an integrated *egl-5::gfp* reporter (Ferreira et al., 1999) and checked for derepression in individual cells after knockdown of the LSM2-8 complex, comparing expression in adult males treated with mock RNAi and RNAi against *lsm-7* and *mes-2*. As reported in Ross and Zarkower, males lacking *mes-2* activity displayed ectopic derepression of this reporter in the male tail region (Fig. 4A,B) and occasionally displayed anterior expansions of tail structures (Fig. 4C). Following *mes-2* RNAi we also found a few nuclei ectopically expressing this reporter outside the tail/posterior region (Fig. 4D, white arrowheads). Interestingly, knock-down of a functional LSM2-8 complex (*lsm-7* RNAi) led to a very similar significant ectopic derepression of the *egl-5* HOX reporter (Fig. 4A,B) and also occasionally provoked anterior expansions of the tails (data not shown). Quantitation of cells expressing EGL-5::GFP revealed on average 20 fluorescent cells per worm under mock RNAi conditions, and about 45 cells following either *mes-2* or *lsm-7* RNAi (Fig. 4B). Thus, cell-specific HOX locus repression is LSM2-8 controlled, further linking LSM-8 to the Polycomb pathway.

**Figure 4:**
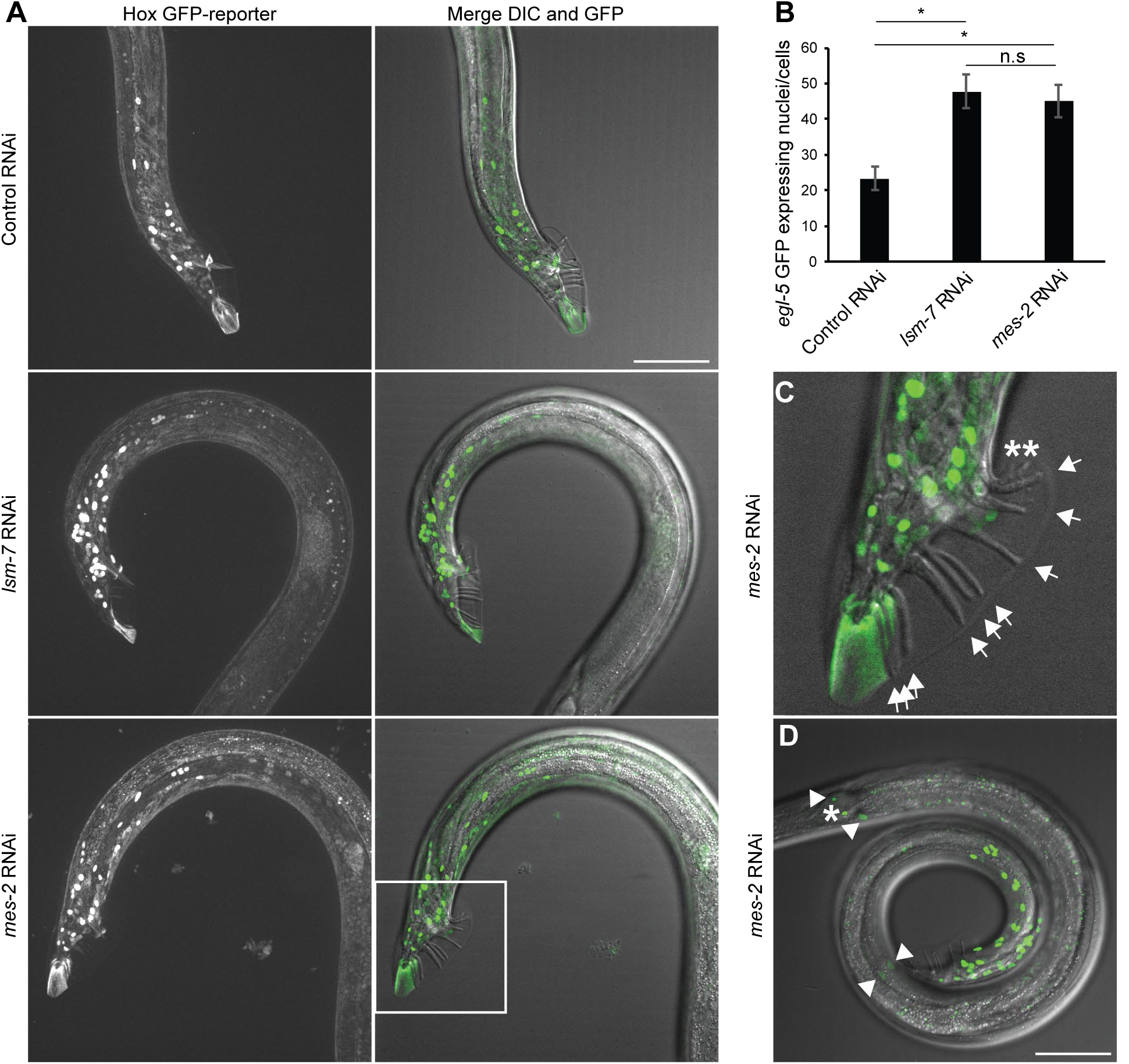
*lsm-7*, like the EZH2 homolog *mes-2*, is required to silence the *egl-5* Hox gene. **A,** On the left, Z-projection of confocal images showing the GFP fluorescence of the *egl-5* GFP HOX reporter (*bxIs13*) under Control (mock/L4440) RNAi, *lsm-7* and *mes-2* (EZH2 homolog) RNAi conditions, in adult males. On the right, merged images of the Z projection of the GFP signal with the DIC image at the best focal plan to visualize the rays of the male tail. Bar, 50 μm. **B,** Quantification of the number of expressing *egl-5* GFP nuclei/cells under the indicated RNAi conditions per proximal region of worms. Student’s t test p-values: *<0.001; n.s = p> 0. 5. N=3; n= 18, 19, 17, bars = s.e.m. **C,** Enlarged male tail inset as in (A) showing the 9 normal rays by arrows and an example of 2 ectopic abnormal rays in *mes-2* (asterisks). **D,** *egl-5* GFP derepression is observable mostly in male tail region, as in (A) but a few nuclei (0 up to 4, as shown by the arrowheads) could also exhibit this derepression in other regions of the worm in *lsm-7* and *mes-2* RNAi conditions. The nucleus indicated by an asterisk express *egl-5* GFP in all conditions tested. Bar, 50 μm.

### *lsm-8* mutation does not induce transcription from both strands nor alter splicing efficiency

To understand the mechanism of LSM2-8 silencing, we carried out a careful analysis of strand-specificity by mapping the RNAs recovered in the *lsm-8* mutant. This showed that derepression occurs exactly over normal gene-coding sequences, without a detectable increase of inaccurate termination or initiation; nor did we detect transcripts from the complementary strand (Fig. S5A). Given that the LSM2-8 complex is known to stabilize and bind U6 snRNA in yeast and plants (Beggs, 2005; Perea-Resa et al., 2012; Zhou et al., 2014) (Fig. 1G), and that the LSM2-8 complex co-precipitates both with U6 snRNA (Fig. S6A) and with a factor involved in U6 snRNA stability (Ruegger et al., 2015), we examined the RNA-seq data for splicing defects. Surprisingly, comparison of exon-exon junction reads in *lsm-8 vs* WT RNA-seq revealed no prominent changes in splicing events, and notably, no intron retention (Fig. S6B). Indeed, out of 134’836 splicing junctions examined, only 18 exon-exon junctions, which mapped to 13 genes, were reproducibly affected (<0.02%; Fig. S6B); none of these was an *lsm-8^-/-^* upregulated gene. Accurate intron splicing is illustrated in Fig. S5. This makes it very unlikely that altered splicing is the source of *lsm-8*^-/-^ mediated derepression.

The RNA-IP done under native conditions confirmed that LSM2-8 complex co-precipitates with U6 snRNA in *C. elegans* (Fig. S6A), as in other species, and further showed that LSM2-8 can associate with a transcript it regulates. There was no significant association of LSM2-8 with a control transcript that is not regulated by LSM2-8 (Fig. S6A).

### LSM2-8 silences gene expression cooperatively with XRN-2

We next examined potential cofactors of LSM2-8 that might contribute to mRNA level regulation. The first candidate was XRN-2, which we showed could derepress the heterochromatic reporter (Fig. 2), and which has been proposed to work with LSM2-8 in budding yeast (Tharun, 2009). We monitored its role in silencing genomic loci by comparing RNA-seq data from *lsm-8^-/-^* L3 larvae with an existing transcriptome of WT L4 larvae treated with *xrn-2* RNAi (Miki et al., 2016) (Fig. 5A). 71% of the genes upregulated by *lsm-8^-/-^* were also upregulated by *xrn-2* RNAi (Fig. 5A, yellow) and 95% of those genes are enriched for H3K27me3 (Tables S32-S3). This argues that LSM-8 and XRN-2 likely function on the same pathway with respect to heterochromatin silencing. Nonetheless, a subset of LSM2-8 target genes (< 33%, in pink) were unaffected by down-regulation of XRN-2. It is unclear if those reflect experimental differences or a subset of genes silenced by LSM2-8 through another mechanism. As expected, more genes were affected by *xrn-2* RNAi than by the *lsm-8^-/-^* mutation (green), given that XRN-2 is involved in a number of other RNA processing events (Miki and Grosshans, 2013; Miki et al., 2014). Consistent with the fact that XRN-2 has broader functions, *xrn-2* deletion leads to early larval arrest (Miki and Grosshans, 2013; Miki et al., 2014), which is a more severe phenotype than that observed in *lsm-8* mutants (Fig. 2; Fig. S2). Interestingly, we found that genes silenced by *xrn-2* independently of *lsm-8* (green) were not enriched for H3K27me3 (Fig. S4D), which suggests that XRN-2 itself is not specific for RNA from Polycomb-marked genes, while the LSM2-8 complex is.

**Figure 5:**
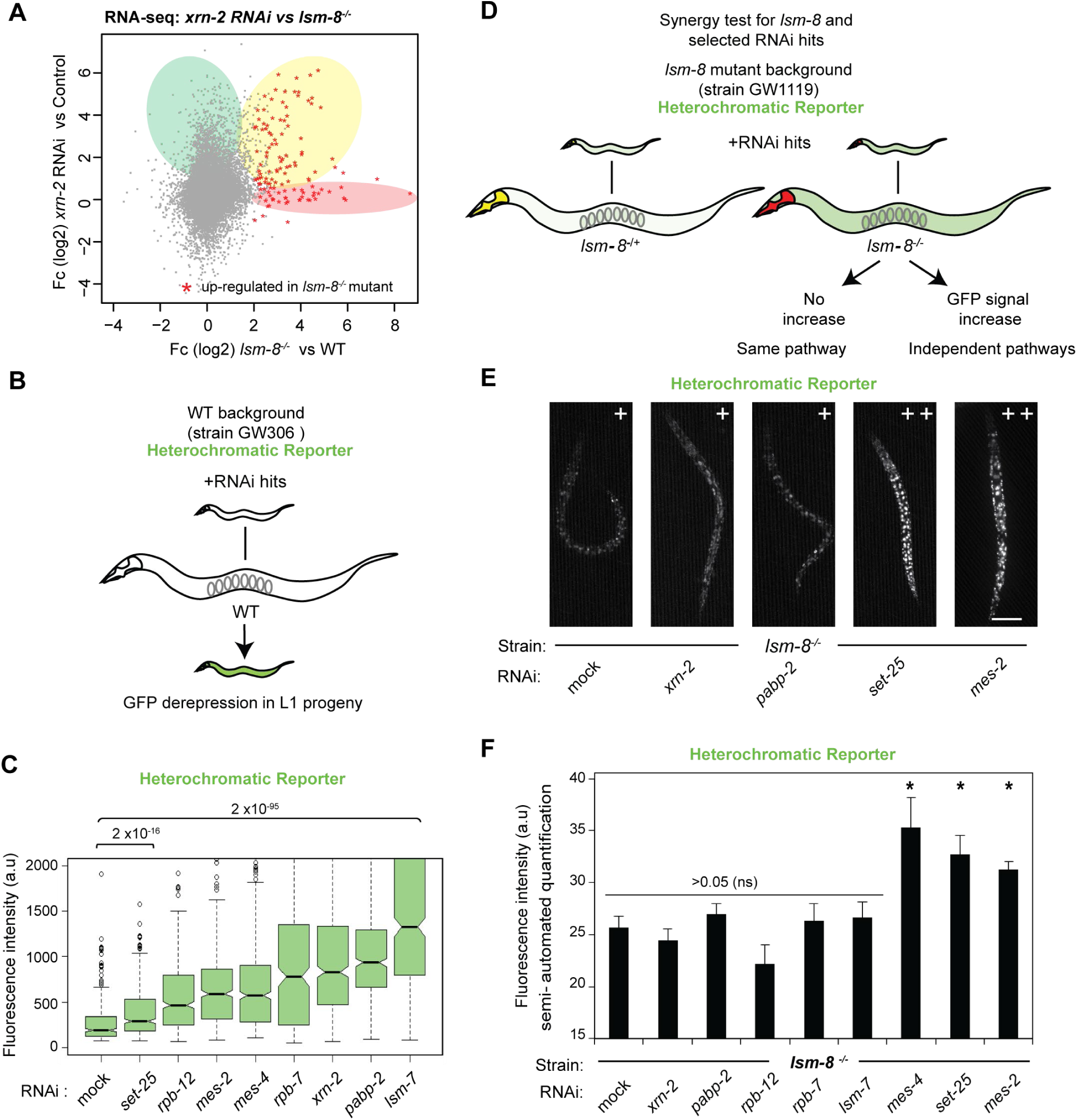
LSM2-8 and XRN-2 work on the same silencing pathway. **A,** Scatter plot comparing relative gene expression changes of *lsm-8^-/-^* L3 larvae (this study) and *xrn-2* RNAi treated L4 (Miki et al., 2016). Common upregulated genes are shaded yellow; 71% of genes upregulated in the *lsm-8* mutant (FDR <0.05 and Fc >4) are also upregulated to some extent (50% increase) in *xrn-2* depleted worms. *lsm-8^-/-^*-specific upregulated genes are shaded pink; *xrn-2* RNAi-specific are shaded green. **B,** Experimental flow for testing the involvement of additional factors in LSM-8 mediated silencing. RNAi experiments were performed in parallel in WT (B) and *lsm-8* mutant (D) backgrounds, from strains GW306 and GW1119, respectively, both carrying the same heterochromatic reporter *pkIs1582*. Derepression assay in WT background confirming derepression following RNAi of indicated factors and RNAi efficiency. **C,** Quantitation of derepression as GFP intensity from the heterochromatic reporter *pkIs1582* in strain GW306, scored in L1 progeny under different RNAi conditions. Fluorescence intensities are displayed as notched box plot. Relevant p values (t-test) are indicated above. n= 500, 500, 500, 500, 500, 500, 295, 500, 500, respectively. **D,** Scheme for analysis of epistasis of RNAi targets with *lsm-8* deletion in larvae progeny of heterozygotic *lsm-8+/-* worms bearing the reporter *pkIs1582*. **E,** Fluorescence microscopy of L4 larvae showing same/ non-additive (+) and additive (++) derepression of the reporter *pkIs1582* in *lsm-8^-/-^* worms under indicated RNAi conditions. Bar, 50 μm. **F,** Quantitation of GFP intensity by semi-automated analysis of microscopic images of the reporter *pkIs1582* in *lsm-8^-/-^* worms under indicated RNAi conditions. *: p<0.005, bars = s.e.m N=2, n = 55, 45, 22, 11, 10, 23, 25, 25, 85.

To identify additional cofactors that might cooperate with LSM2-8 and XRN-2, we concentrated on factors involved in RNA transcription or processing. Based on earlier work showing that RNA Pol II subunits cooperate with budding yeast LSM1-7 in cytoplasmic RNA decay (Haimovich et al., 2013; Lotan et al., 2007), we examined the effects of two RNA Pol II subunits (*rpb-12, rpb-7*) and the type II poly(A) binding protein *pabp-2* (*Hs*PABPN1 and *Sp*Pab2). We find that like *xrn-2* RNAi, *rpb-12, rpb-7* and *pabp-2* RNAi’s derepress the heterochromatic reporter on their own (Fig. 5B,C). This is not true for all genes implicated in RNA metabolism, as exemplified by *cgh-1* (*Hs*DDX6 and *Cs*Dhh1) and *pab-2* (*Hs*PABPC1 and *Cs*Pab1), two factors involved in RNA regulation that have the same effect as mock RNAi (Fig. 5C). This suggested that RPB-12, RPB-7 and PABP-2 may also contribute to silencing the heterochromatic reporter.

To see if these genes act on a common pathway with LSM-8, we performed RNAi against these factors both in a WT and in the *lsm-8^-/-^* strain, and scored whether their effects were additive or epistatic with the loss of LSM-8. The *lsm-8* mutant worms were treated with RNAi and scored for GFP derepression in the homozygous *lsm-8^-/-^* (red pharynx) worms of the next generation (Fig. 5D). We found the down-regulation of *xrn-2, pabp-2 rpb-12* or *rpb-7,* was completely epistatic with *lsm-8* deficiency for reporter derepression, as was the *lsm-7* RNAi control (Fig. 5E,F). RNAi against the Polycomb HMT *mes-2* was additive with *lsm-8* deletion, albeit less so than either *set-25* (H3K9me3 HMT) or *mes-4* (H3K36 HMT). We conclude that LSM2-8 acts on a pathway of silencing that is dependent on XRN-2-mediated RNA metabolism. The fact that *lsm-8* and *mes-2* are not fully epistatic argues that Polycomb-mediated silencing does not depend entirely on LSM-8. This is to be expected, as PRC2 complexes can repress active transcription, while LSM2-8 likely acts post-transcriptionally (Fig. 5E,F; see below).

Given the striking correlation of LSM-8-sensitivity with H3K27me3 (Fig. 3), we next examined if LSM2-8 silencing requires the presence of H3K27me3. To this end, we tried to combine the *mes-2* mutant with the balanced *lsm-8* deletion, but as both mutations provoke sterility, this was not possible. Moreover, RNAi was extremely inefficient in the *mes-2* null background for all clones tested (e.g. *lsm-7, ubq-1, let-607*). Therefore, we asked instead whether the loss of LSM2-8 alters the accumulation of H3K27me3 on affected genes. Indeed, quantitative ChIP-qPCR for H3K27me3 on *lsm-8* upregulated genes had a significant decrease (>50%) in H3K27me3 levels in *lsm-8* vs WT larvae (Fig. 6A), while *lsm-8*-insensitive genes did not. This suggests that the LSM2-8 complex feeds back to maintain H3K27me3 levels, either directly or indirectly, specifically at the H3K27me3-marked loci that are sensitive to *lsm* ablation.

**Figure 6:**
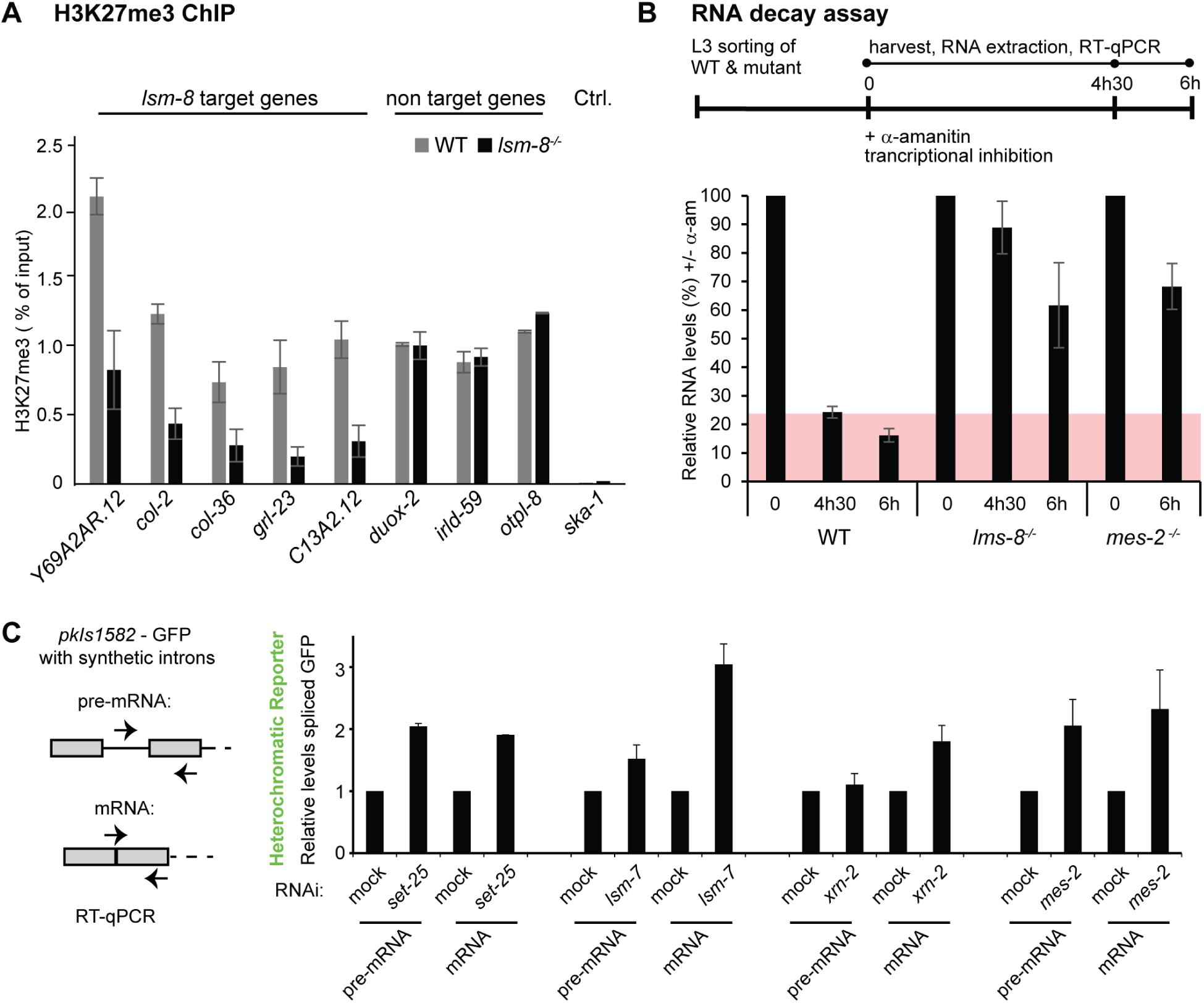
LSM2-8 mediates silencing primarily through RNA degradation. **A,** H3K27me3 ChIP-qPCR on target genes in WT and *lsm-8^-/-^* worms. Three categories of genes were assessed. Genes that are upregulated in *lsm-8^-/-^* worms and enriched for H3K27me3 mark in WT (*lsm-8* target genes), genes that are enriched for the H3K27me3 mark in WT but do not change in expression in the *lsm-8* mutant (non-target genes), and the control *ska-1*, a gene not enriched for H3K27me3 in WT and with no expression change (ctrl). N=3, n=3, bars = s.e.m. **B,** WT *lsm-8^-/-^* and *mes-2^-/-^* worms treated with 50 µg/ml of the transcriptional inhibitor α-amanitin for indicated times. Levels of transcripts from 3 genes regulated by LSM-8 or two that are not (see individual genes and results in Fig. S7) were tested by RT-qPCR and normalized to 18S rRNA levels. 0h was defined as 100%. N=3, n=3, bars: s.e.m. **C,** RNA levels of the pre-mRNA and mRNA of GFP from the heterochromatic reporter *pkIs1582* from the strain GW306 were determined by RT-qPCR and normalized to *pmp-3* mRNA. The levels on mock RNAi conditions are defined as 1. N = 2, 3, 2, 3, respectively; n = 3, bars = s.e.m. *mes-2* RNAi depletes MES-2/PRC2-like and H3K27me3 levels; *set-25* RNAi depletes SET-25 and H3K9me3 levels.

### LSM-8 and XRN-2 cooperate to promote RNA decay

The partial feedback on H3K27me3 levels by the LSM2-8 complex on genes it regulates is unlikely to account for the strong *lsm-8^-/-^* impact on mRNA level, thus we next tested whether the LSM2-8 complex influences mRNA turnover rates. In WT, *lsm-8 and mes-2* mutant backgrounds we added α-amanitin, an inhibitor of RNA pol II and pol III elongation, to L3 stage larvae, and monitored RNA decay over 6 hours by RT-qPCR. The stability of mRNAs from genes known to be sensitive to *lsm-8^-/-^* (Table S2), was compared between the *lsm-8* mutant, a *mes-2* mutant, and WT L3 larvae, normalizing mRNA levels to the 18S rRNA, as its synthesis is insensitive to α-amanitin (Fig. S7A). The *lsm-8* sensitive genes do show a delayed rate of decay in the absence of LSM-8 (Fig. 6B). The rate varied slightly among the three genes monitored (*far-3, grl-23* and *ZK970.2*, Fig. S7), yet all were significantly different from two control genes (*eft-3*, *F08G2.8*), which were unaffected by the *lsm-8* mutation (Fig. S7B). This indicates that the LSM2-8 complex is required to promote the degradation of the transcripts it regulates. Similarly, transcript levels from the *lsm-8*-sensitive genes *far-3* and *ZK970.2* were strongly upregulated in *xrn-2* RNAi-treated worms, with a log2 fold change of 4.9 and 4.0, respectively. This suggests that the elevated levels of mRNA detected in *lsm-8^-/-^* worms stem from RNA stabilization. Importantly, by monitoring RNA decay in the homozygous *mes-2* mutant we scored a similar increase in mRNA stability as that found in the *lsm-8* mutant, for transcripts sensitive to LSM-8 ablation (Fig. 6B and S7B). This suggests that H3K27me3, or at least MES-2 is required for the LSM8-mediated RNA degradation.

To see whether RNA degradation primarily acts on nascent transcripts (i.e., is co-transcriptional) or occurs post-transcriptionally on mature mRNA, we compared the newly transcribed pre-mRNA levels with those of spliced mRNAs derived from the heterochromatic reporter *pkIS1582*. We scored the levels of unspliced and spliced mRNA following *lsm-7*, *xrn-2* and *mes-2* RNAi, and used *set-25* RNAi as a control (Fig. 6C). If repression or degradation occurs at the level of transcription, we expect the increase in pre-mRNA and in mRNA to be equal. If spliced mRNA levels are higher than the pre-mRNA levels following RNAi, then the drop in mRNA level is likely to be a post-transcriptional event. Quantitative PCR showed that the loss of LSM-7 and XRN-2 showed a much stronger accumulation of mature mRNA over pre-mRNA (Fig. 6C), whereas the loss of the H3K9me HMT SET-25 affected pre-RNA and mRNA levels equally (Fig. 6C). This is consistent with SET-25 playing a role in transcriptional repression, while LSM2-8 and XRN-2 appear to act primarily on mature RNAs. The down-regulation of *mes-2* had an intermediate effect between *set-25* and *lsm-7* RNAi, consistent with roles in both the targeting of LSM2-8 and reducing transcription (Fig. 6C).

Taken together our data suggest that LSM2-8/XRN-2 is essential for the post-transcriptional degradation of RNAs originating from Polycomb-marked genes. We conclude that facultative heterochromatic silencing is achieved in *C. elegans* on two levels: one reflects a reduction in transcriptional efficiency, while the second is primarily post-transcriptional, requiring LSM2-8 and XRN-2 to degrade nuclear RNAs prior to cytoplasmic export (Fig. 7).

**Figure 7:**
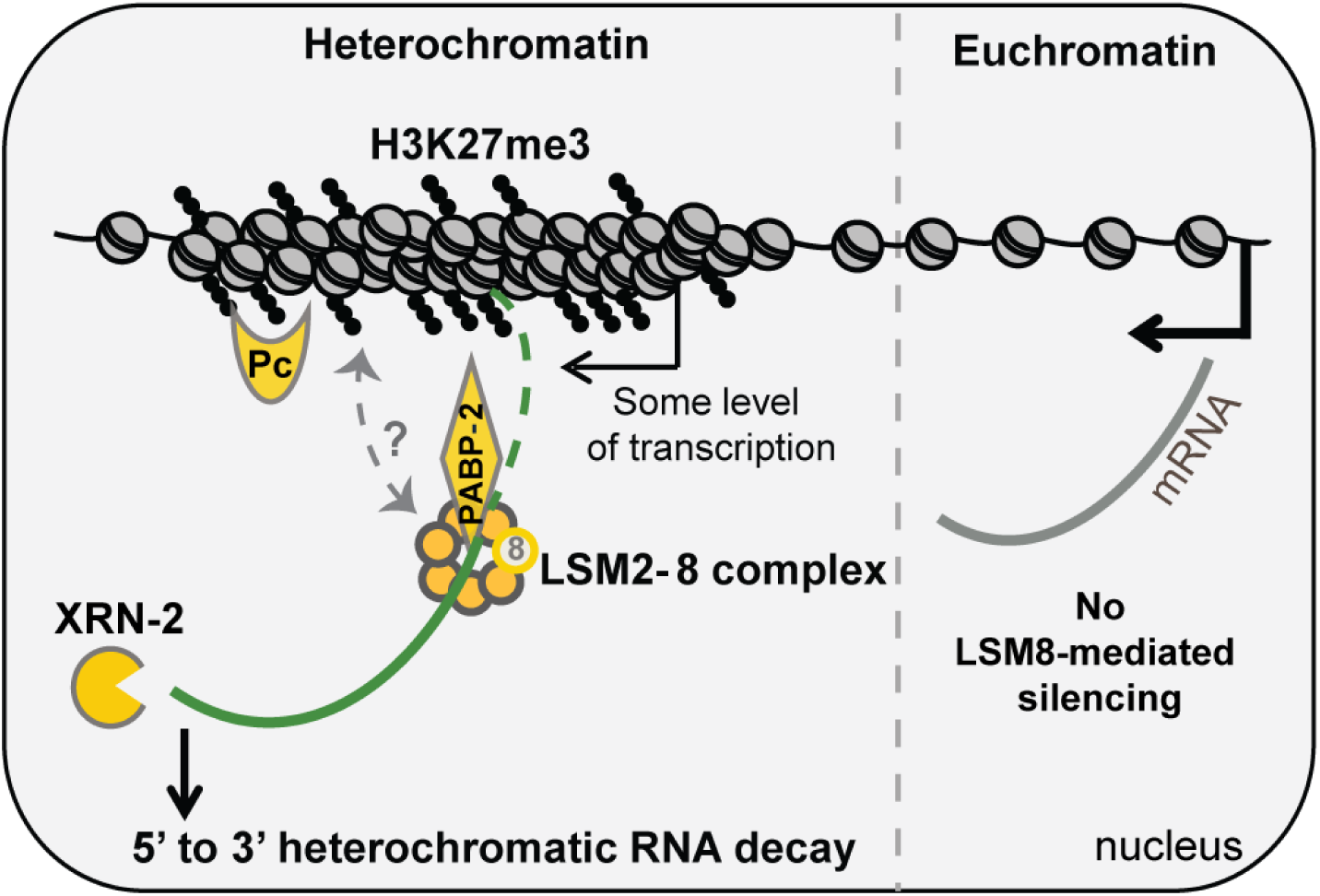
LSM2-8 complex and XRN-2 silence transcripts arising from heterochromatic H3K27me3-enriched domains through RNA degradation. The LSM-8 mediated silencing pathway makes use of XRN-2 ribonuclease, and may involve other transcript binding factors, such as PABP-2 (*Hs*PABPN1, see Discussion). We hypothesize that RNA arising from H3K27me3 genomic regions that are controlled by the LSM2-8 complex may acquire a specific feature during transcription (*e.g.* a specific structure, RNA modification, 3’UTR, poly-A/U tail, or specific RNA binding protein(s)), that allows recognition and processing by LSM2-8. LSM2-8-mediated silencing also feeds back to regulate H3K27me3 levels on LSM-8-regulated genes, although it is unclear if the interaction with PRC2 or H3K27me3 is direct (dotted arrow). The LSM-2-8-mediated silencing of H3K27me3-bound loci defines a selective post-/co-transcriptional silencing through RNA decay, beyond the transcriptional repression attributed to facultative heterochromatin.

## Discussion

We have shown in nematodes that a conserved nuclear complex, LSM2-8, contributes to the selective repression of heterochromatic reporters and of a subset of genomic loci bearing the H3K27me3 epigenetic mark, through the post-transcriptional degradation of mRNA. In contrast, a related cytoplasmic complex, LSM1-7, appears to have no role in heterochromatic silencing. Unlike heterochromatin-linked RNA processing pathways in plants and fission yeast, which include the RITS, TRAMP and exosome complexes (Buhler, 2009; Coy and Vasiljeva, 2011; Grewal and Elgin, 2007; Moazed, 2009), the silencing mediated by LSM-2-8 does not target H3K9me3-marked genes specifically and the complex acts independently of the two *C. elegans* H3K9-specific HMTs. Our data show instead that the LSM2-8 complex specifically reduces the stability of transcripts arising from H3K27me3-tagged genes, mediating their decay by the 5’-3’ exoribonuclease XRN-2 (Fig. 7). Genetically, we also implicate the RNA pol II factors RPB-7 and RPB-12, and the type II poly(A) binding protein (PABP-2) in the silencing of the heterochromatic reporter. Importantly, LSM2-8-mediated silencing of endogenous loci occurs almost exclusively at genes carrying H3K27me3, as revealed by transcriptomes of mutant L1 and L3 larval stages. Furthermore, as shown by the RNA decay assay in the *mes-2* mutant (Fig. 6), it seems to require the presence of MES-2 (the *C. elegans* EZH2 homolog). Given that derepression of the heterochromatic reporter occurred in all somatic tissues of L4 larvae, we argue that LSM2-8-mediated silencing is likely to be broadly relevant, because reporter derepression could be monitored in all somatic cells at all stages of development.

Figure 7 illustrates the proposed mode of action, whereby the LSM2-8 complex is targeted either through MES-2 itself, through an unknown ligand of H3K27me3 or through characteristic aspects of the nascent transcripts (e.g. poly-U stretches) to mediate post-transcriptional degradation by XRN-2. This silencing acts in parallel to H3K27me3-mediated transcriptional repression, apparently increasing the robustness to the Polycomb-mediated repression of developmental genes. The conserved nature of the LSM proteins and of the other factors implicated in the LSM2-8 silencing pathway (XRN-2, MES-2, PABP-2, etc) suggests that this mechanism may be active in other species.

Based on RNAi and genetic epistasis studies, we propose that the RNA decay mediated pathway identified here may involve the type II poly(A) binding protein (PABP-2, (Hurschler et al., 2011) and the RNA Pol II subunits, RPB-12 and RBP-7, although the pathway is independent of DCAP-2, XRN-1, LSM-1 (Fig. 1H) and HPL-2 (data not shown). PABP-2 (*Hs*PABPN1 and *Sp*Pab2) is particularly interesting, because PABP-2 is nuclear and appears to regulate 3’UTR and poly(A) tail length (Kuhn et al., 2009). Moreover, it binds nascent RNAs early during the elongation step (Beaulieu et al., 2012; Lemieux and Bachand, 2009). Given that the LSM2-8 complex is known to bind to the 3’ oligo(U) tail of the U6snRNA (Zhou et al., 2014), as well as the 3’ poly(A+) tail of nuclear RNAs (Kufel et al., 2004), we hypothesize that PABP-2 may regulate LSM2-8 specificity by modulating the 3’end of mRNAs at H3K27me3-marked domains. In that way, PABP-2 might serve as a “mediator”, to link the chromatin state to the transcripts targeted by LSM2-8.

It is unclear whether any molecular criteria other than H3K27me3 modification are necessary to target mRNAs for derepression by LSM2-8/XRN-2. We note that the fraction of H3K27me3-marked genes that is sensitive to LSM2-8 repression is likely to be higher than that detected by RNAseq of whole larvae. First, if we apply a 1.5-fold change cut-off, instead of >4-fold for steady-state mRNA levels in L3 larvae, we find 21% of all Polycomb-marked genes are upregulated by loss of LSM-8. This is a highly significant fraction, given the fact that the removal of a silencing pathway is usually insufficient to trigger gene expression: a promoter must also be bound by relevant transcription factors to drive RNA PolII-dependent transcription. A second factor that has limited our ability to score all *lsm-8* sensitive transcripts is the fact that our genomic transcriptome analyses had to be done on RNA extracted from whole animals, due to the lethal nature of the *lsm-8* deletion. Transcripts that are upregulated in a restricted number of cells, or in a specific cell-type, can easily be missed in whole animal RNA libraries, and require detection by sensitive imaging methods that can monitor derepression in a cell-specific manner. This was implemented for the HOX gene *egl-5* (Fig. 4), a Polycomb target that is sensitive to *lsm-8* ablation, even though its upregulation occurred only in ∼45 posterior cells in males (Bender et al., 2004; Hench et al., 2015; Ross and Zarkower, 2003; Soshnikova and Duboule, 2009; Yuzyuk et al., 2009). A similar pattern of *egl-5* derepression was scored upon loss of MES-2, the EZH2 homologue, reinforcing the link of LSM2-8 and XRN-2 to Polycomb (Fig. 4, Table S2). From this result, we infer that the extent to which LSM2-8 and XRN-2 regulate developmentally relevant transcripts from H3K27me3-marked loci is likely be more extensive than what whole animal RNA-seq can reveal.

How LSM2-8 recognizes H3K27me3-marked genes is unclear. While LSM proteins do not contain methyl lysine-binding chromodomains, they may recognize specifically modified RNAs or short ribonucleotide motifs. The recognition of RNAs arising from H3K27me3-marked domains thus could entail features acquired during transcription, such as a specific secondary structure, an RNA modification, a modified or alternative caps, variant poly-A or U tails, or may bind a unique RNA-binding factor. We do not yet rule out that there are H3K27me3-binding chromodomain proteins that help target the LSM2-8 complex, although in this case they must act redundantly. Examples of redundant H3K27me3 binding factors include two recently identified H3K27me3 ligands, CEC-1 and CEC-6 (Saltzman et al., 2018), whose loss together with ablation of an H3K9me-binding protein, CEC-3, caused a mortal germline phenotype. We note, however, that unlike the *LSM* genes, RNAi against these chromodomain proteins did not derepress heterochromatin in a genome-wide screen (Towbin et al., 2012), nor in a targeted screen that monitored all *C. elegans* methyl lysine-binding factors by RNAi (Gonzalez-Sandoval et al., 2015). Thus, if they are involved in the LSM2-8 pathway, it must be in a redundant fashion.

PRC1 subunits are not well conserved in *C. elegans* (Wenzel et al., 2011), however two *C. elegans* Polycomb-like factors, SOR-1 and SOP-2, have also been shown to be essential to prevent ectopic HOX gene activation and have RNA binding activity (Zhang et al., 2004; Zhang et al., 2006). Such factors might also provide a link between Polycomb targets and the RNA degradation machinery.

We note that RNAi against RNA Pol II subunits RBP-7 and RBP-12 derepressed the heterochromatic array in a manner epistatic with *lsm-8* deficiency. Intriguingly, in *S. pombe* the RBP-7 homolog has been implicated in centromeric repeat transcription and RNAi-directed silencing (Djupedal et al., 2005), while in *S. cerevisiae,* the same RNA Pol II subunit plays a direct role in Pat1/Lsm1-7 mediated mRNA decay in the cytoplasm (Haimovich et al., 2013; Lotan et al., 2007). It is not yet clear what links these various observations, but it should not be excluded that RPB-7 and RPB-12 subunits might mark LSM2-8 regulated transcripts to signal degradation by XRN-2, given their epistatic effect on reporter derepression. A systematic conditional screen of all RNA Pol II subunits and their relationship to *lsm-8* will be needed to clarify if RNA Pol II subunits participate directly in the silencing pathway described here.

We found no involvement of the decapping enzyme, DCAP-2, or of CGH-1, the closest homolog in *C.elegans* of the yeast decapping enhancer Dhh1 (Nissan et al., 2010) in the LSM2-8 pathway of degradation, although it is reasonable to expect that a nuclear decapping function might be needed to sensitize target RNAs to degradation by XRN-2. We note that XRN-2 has multiple nuclear functions (Miki and Grosshans, 2013), yet we did not see evidence for aberrant termination events in *lsm-8* deficient worms, nor are *lsm-8* sensitive genes enriched for loci that require XRN-2 for termination (Miki et al., 2017).

Importantly, we document here by ChIP a drop in H3K27me3 on *lsm-8*-sensitive genes in *lsm-8*^-/-^ animals, suggesting that the LSM2-8-mediated RNA degradation pathway feeds back to maintain H3K27me3 levels. Interestingly, a recent but still debated suggestion was made that ncRNAs, such as Xist or HOTAIR, or other PRC2 binding RNAs, may help target Polycomb in *cis* or in *trans* to target genes (Brockdorff, 2013; Johnson and Straight, 2017; Ringrose, 2017). Understanding how this might relate to the mechanism described here, where an RNA binding and degradation complex contributes to gene silencing and H3K27me3 maintenance in worms, is a topic of future research. It is nonetheless highly significant that epigenetic information (H3K27me3) is being sensed by an RNA binding complex, that in turn seems to reinforce the chromatin state (Fig. 7). Thus, despite the fact that LSM2-8 works primarily post-transcriptionally (Fig. 6), the pathway may feed back to stabilize the mark that characterized the source of the transcript it degrades.

Overall, our study shows that facultative heterochromatin silencing in a multicellular organism makes use of a mechanism of selective transcript degradation, and not only of transcriptional repression. LSM2-8-mediated gene silencing furthermore links a specific epigenetic state to transcript degradation, adding an additional layer of control over differentiation and development.

## Acknowledgements

The accession number for the RNA-seq data is NCBI Gene Expression Omnibus is GSE92851. Some strains were provided by the *Caenorhabditis* Genetics Center (CGC), which is funded by NIH Office of Research Infrastructure Programs (P40 OD010440). We thank I. Katiç, the FMI Genomics and Microscopy facilities, P. Zeller and M. Fukushima for technical help, advice and discussion, and T.S. Miki for access to data and discussions. We thank M. Bühler, H. Großhans and W. Filipowicz for discussions and proofreading of the text. The authors acknowledge support of the Novartis Research Foundation, as well as a Marie Curie Intra-European grant (#PIEF-GA-2010-276589) and Swiss National Science Foundation Marie-Heim Vögtlin grant (#PMPDP3_151381, # PMPDP3_168717) to AM; SNF grant (#310030B_156936) to SMG and support of the NCCR RNA & Disease to H. Großhans (to FA.).

## Author Contributions

A.M. planned and executed most experiments, evaluated results and wrote the paper; S.M.G. planned experiments, evaluated results and wrote the paper; D.G. analyzed with A.M. the RNA-seq data and other genome wide data, C.S analyzed the L1 RNA-seq data, J.P. performed the H3K27me3 ChIP-qPCR experiment and analysis; V.K. performed the gonad staining and analysis and F.A provided help for the *lsm-8* mutant strain generation.

## Author Information

Affiliation: Friedrich Miescher Institute for Biomedical Research, Basel, Switzerland. Anna Mattout, Dimosthenis Gaidatzis, Jan Padeken, Christoph Schmid, Florian Aeschlimann, Véronique Kalck and Susan M. Gasser

Faculty of Natural Sciences, University of Basel, Basel, Switzerland. Florian Aeschlimann, Susan M Gasser

Université Paul Sabatier-CNRS UMR 5088, Toulouse France. Anna Mattout Competing financial interests: The authors declare no competing financial interests. Corresponding Author: Correspondence to Susan M. Gasser

## Supplementary Information

**Figure S1:**
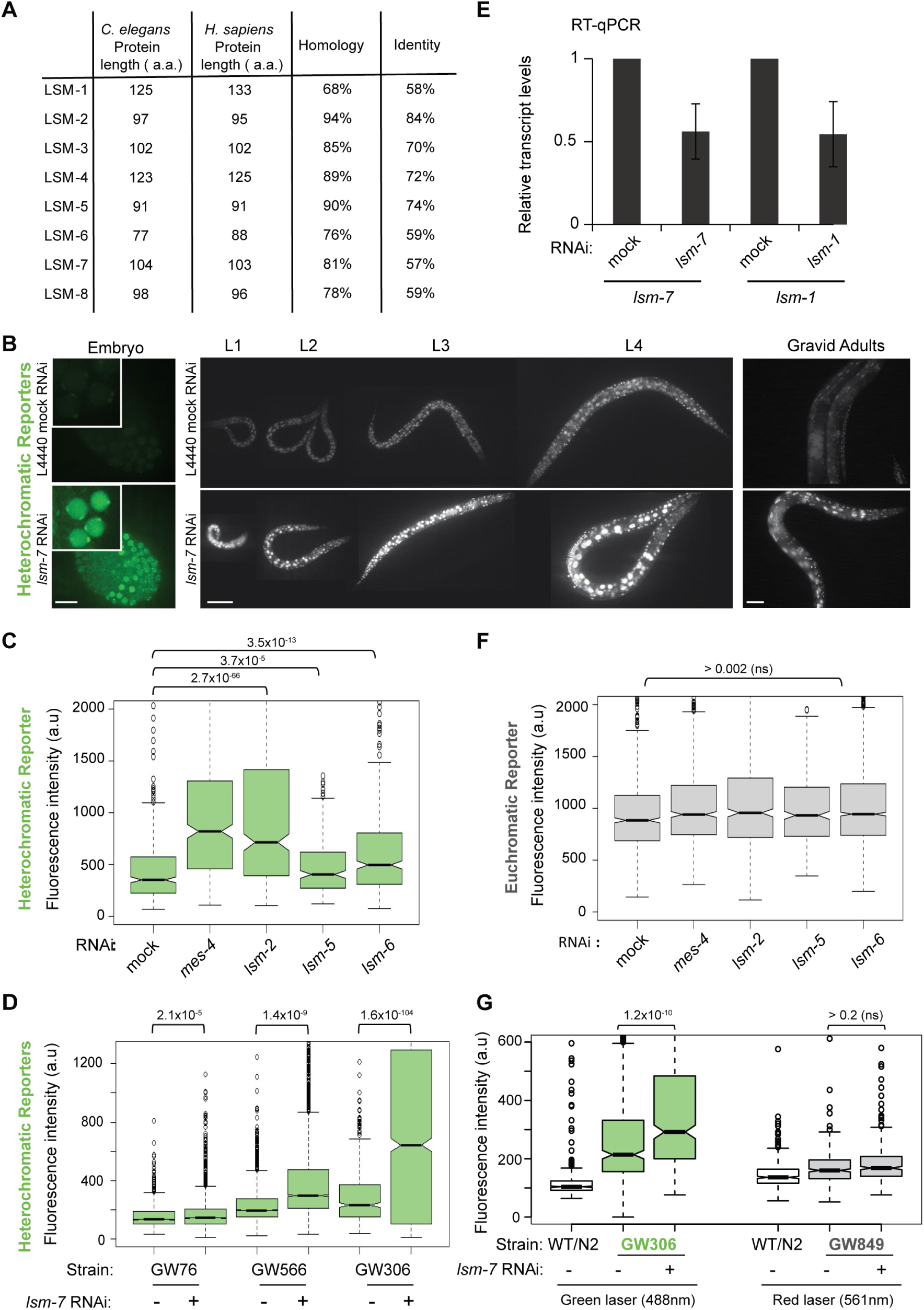
LSM proteins are highly conserved and silence heterochromatic, but not euchromatic reporters. **A,** LSM protein length and conservation between *C. elegans* and *H. sapiens*. **B,** Heterochromatic reporters derepression at all developmental stages. The derepression (GFP live imaging) under *lsm-7* RNAi compared to the control RNAi condition is shown for the embryonic stage (heterochromatic reporter from strain GW566, Table S1). Nuclei are enlarged in the inset. Bar, 10 μm. The derepression is also shown for all larval stages L1-L4, bar: 50 μm and gravid adults, bar: 100 μm (heterochromatic reporter from strain GW306, Table S1). **C,** Quantitation of derepression assays. In L1 progeny under *gut-2/lsm-2, lsm-5, lsm-6* and control RNAi conditions (mock: negative control and *mes-4*: positive control), the GFP fluorescence intensity of the heterochromatic reporter *pkIS1582* from the strainGW306 was measured by the worm sorter. F2: second generation. n= 375 for each condition. The GFP fluorescence intensities are displayed as in Fig. 1. **D,** Quantitation of derepression of different heterochromatic reporters. Fluorescence intensities are displayed as notched box plot in arbitrary units (a.u), whiskers = 1st and 3rd quartiles, black lines: median, black circles outliers. P-values are indicated and were calculated in this and all other plots by pair-wise comparisons with the Student’s t-test. In all cases there is a statistically significant reporter derepression upon *lsm-7* knockdown. n= 1460, 2399, 2631, 3850, 634, 1855. **E,** Confirmation of *lsm-1* and *lsm-7* knockdown by RNAi. qPCR analysis of *lsm-7* and *lsm-1* mRNA in L1 worms upon mock, *lsm-7* or *lsm-1* RNAi treatments, done in parallel to the GFP derepression assay. *lsm-7* and *lsm-1* mRNA were normalized to *his-56* and *its-1* mRNA, and values are expressed relative to the levels in mock RNAi condition. N=2, n=3, bars = s.d. **F,** Quantitation of derepression of the euchromatic reporter (GW849) in L1 progeny as in (C). The wmCherry fluorescence intensity (gain 2) of the euchromatic reporter from GW849 strain was quantified. (n.s: not significant). n= 375 for each condition. **G,** Same as in (F), except the same gain settings (gain 1) is used both for the fluorescence of the heterochromatic (GW306) and euchromatic (GW849) reporters. The euchromatic reporter fluorescence is lower in this case than the heterochromatic reporter. n= 370 for all. WT/N2 shows the green and red background fluorescence, respectively, in the absence of the reporter constructs.

**Supplementary Table S1:** List of strains used in this study

| Strain | Original Name / OC | Genotype | Reference |
| --- | --- | --- | --- |
| GW1 | N2* | wild-type, Bristol isolate |  |
| GW76 |  | *gwIs4 [baf-1p::GFP-lacI::let-858 3'UTR; myo-3p::RFP] X* | (Meister et al., 2010) |
| GW306 | NL2507 | *pkIs1582[let-858p::GFP:: let-858 3'UTR; rol-6(su1006)]V* | (Towbin et al., 2012) |
| GW566 |  | *gwIs39 [baf-1p::GFP-LacI::let-858 3'UTR; vit-5p::GFP] III; gwIs4 [baf-1p::GFP-lacI::let-858 3'UTR; myo-3p::RFP] X* | (Towbin et al., 2012) |
| GW653 | YG118 | *ygIs[baf-1p::GFP-lmn-1 Y59C; unc-119(+)] ; unc-119(ed-3)* | (Mattout et al., 2011) |
| GW299 |  | *gwIs25 [tbb-1p::wmCherry-LacI::tbb-2 3'UTR unc-119(+)] unc-119(ed3)* | This study |
| GW886 |  | *ygIs[baf-1p::GFP-lmn-1 Y59C; unc-119(+)] ; set-25(n5021) III. met-2(n4256) III* | This study |
| GW638 |  | *met-2(n4256) set-25(n5021) III* | (Towbin et al., 2012) |
| GW637 |  | *met-2(n4256) set-25(n5021) III; gwIs4 [baf-1p::GFP-lacI::let-858 3'UTR; myo-3p::RFP] X* | (Towbin et al., 2012) |
| GW214 |  | *hpl-2(tm1489) III; gwIs4 [baf-1p::GFP-lacI::let-858 3'UTR; myo-3p::RFP] X* | (Gonzalez et al., 2015) |
| GW468 |  | *mes-2(bn11) unc-4(e120)/mnC1 dpy-10e128() unc-52(e444)II; gwIs4[myo-3::RFP baf-1::GFP lacI let-858] X.* | (Towbin et al., 2012) |
| GW849 |  | *gwSi17 [cec-4p::cec-4::WmCherry::cec-4 3'UTR] II* | (Gonzalez et al., 2015) |
| GW1108 | EG6070 ocx2 | *oxSi221 [eft-3p::GFP + Cbr-unc-119(+)] II (in ttTi5605); unc-119(ed3) III* | (Frøkjær-Jensen et al., 2012) |
| GW638 |  | *met-2(n4256) set-25(n5021) III* | (Towbin et al., 2012) |
| GW1109 | HW1390 ocx4 | *lsm-8 (xe17 [myo2p::mcherry::unc54 3'UTR])IV* | This study |
| GW1125 | ocx6 | *lsm-8 (xe17 [myo2p::mcherry::unc54 3'UTR])IV/nT1[qIs51](IV;V)* | This study |
| GW1119 | ocx6 | *lsm-8 (xe17 [myo2p::mcherry::unc54 3'UTR])IV/nT1[qIs51](IV;V); pkIs1582[let-858::GFP rol-6(su1006)]V* | This study |
| GW1120 | ocx6 | *lsm-8 (xe17 [myo2p::mcherry::unc54 3'UTR])IV ; pkIs1582[let-858::GFP rol-6(su1006)]V* | This study |
| GW1148 | ocx6 | *met-2(n4256) set-25(n5021) III; lsm-8 (xe17 [myo2p::mcherry::unc54 3'UTR])IV/nT1[qIs51](IV;V)* | This study |
| GW923 | VC2663*<br>ocx5 | *lsm-5(ok3431) V/nT1[qIs51](IV;V)* | This study |
| GW925/<br>GW1082 | VC904*<br>ocx5 | *eea-1&gut-2(gk407) V/nT1[qIs51](IV;V)* | This study |
| GW931 | ocx6 | *eea-1&gut-2(gk407) V/nT1[qIs51](IV;V) ; gwIs4 [ myo-3::RFP baf-1::GFP lacI let-858] X* | This study |
| GW933 | ocx6 | *lsm-5(ok3431) V/nT1[qIs51](IV;V); gwIs4 [ myo-3::RFP baf-1::GFP lacI let-858] X* | This study |
| GW1080 | VC2785*<br>ocx5 | *lsm-4&ada-2(ok3151)/mln1[mls14 dpy-10(e128)]II.* | This study |
| GW1004 |  | *gwEx81[WRM062D_B06::gfp::3xFlag]* | This study |
| GW1420 |  | *dcap-2 (tm2470)/ nT1 IV; pkIs1582[let-858::GFP rol-6(su1006)]V* | This study |
| GW1500 |  | *lsm-1(tm3585)/mln1[mls14 dpy-10(e128)]II;*<br>*pkIs1582[let-858::GFP rol-6(su1006)]V* | This study |
| GW1613 | EM599 | *him-5(e1490) V; lin-15B&lin-15A(n765) X; bxIs13* | CGC |

\*CGC: Caenorhabditis Genetics Center
OCx4 : out-crossed 4 times

**Supplementary Figure S2:**
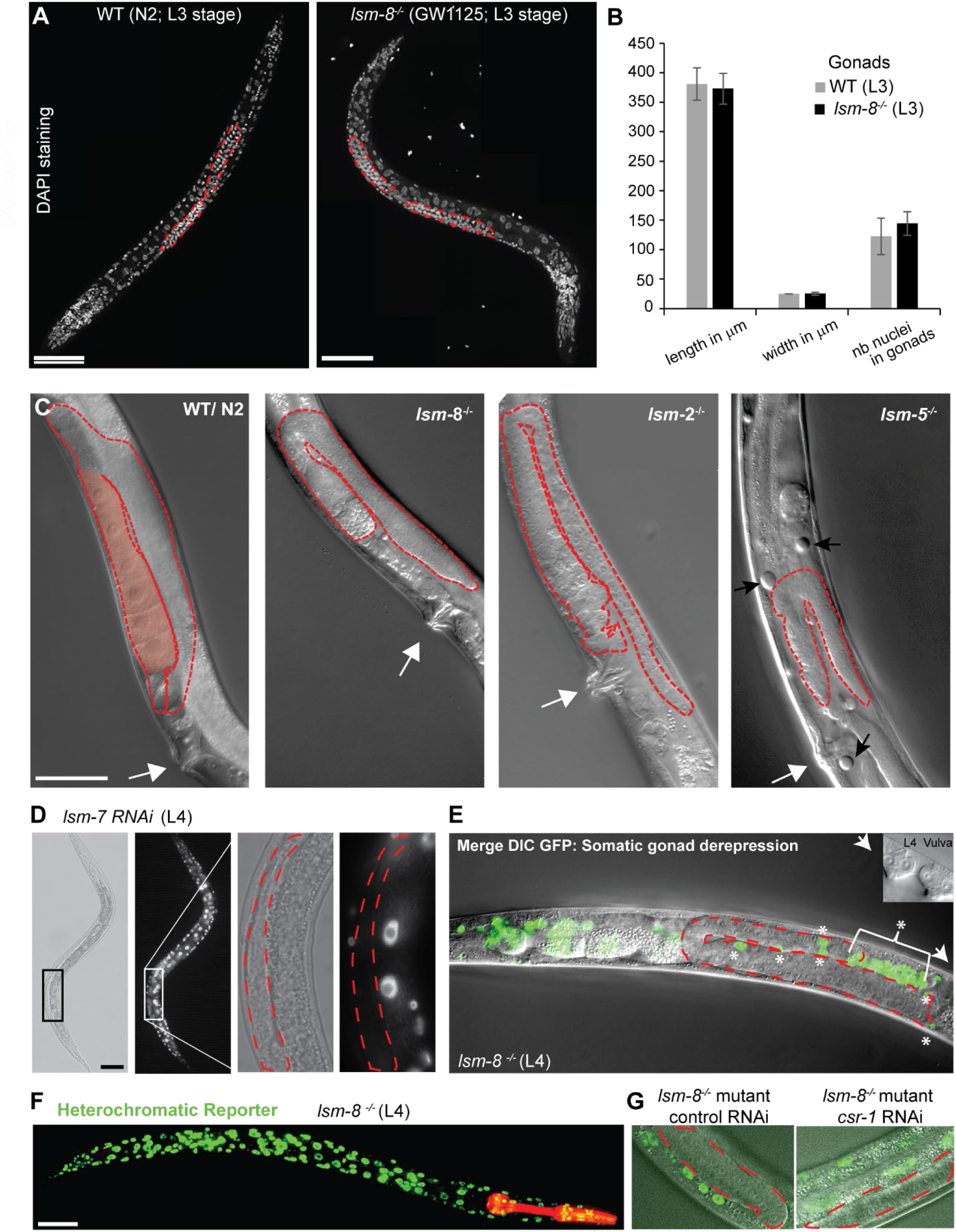
*lsm-8^-/-^* mutant worms are 100% sterile but developing gonads resemble WT through L3 and L4 stages. **A,** Z-projection of confocal images showing fixed DAPI staining of a WT (N2) worm, at L3 stage. Gonad arms are highlighted by the red dashed line and same to right with a *lsm-8^-/-^* L3 larva (GW1125). Bars, 50 μm. **B,** Quantification of the length, width and gonad nuclei count from the DAPI staining of L3 WT and *lsm-8-/-* larvae. n=8. **C,** DIC image of a WT young adult (YA) worm with a normal general anatomy and normal gonad (red dashed line) with oocytes (pink shading). The white arrow indicates the vulva as in YA. DIC image of *lsm-8^-/-^*, *lsm-2^-/-^* and *lsm-5^-/-^* YA worms. The gonad (red dashed line) has no forming oocytes and has an abnormal composition of cells at that stage. Black arrows indicate the presence of vacuoles. Bar, 50 μm. **D,** Heterochromatic reporter (*pkIs1582)* derepression in WT background from strain GW306 following *lsm-7* RNAi in a L4 larva. The enlargement to the right shows the gonad (red dashed line) with germ cells which are not derepressed. Bar, 50 μm. **E,** Merge DIC and live GFP microscopy of *lsm-8* mutant carrying the heterochromatic reporter *pkIs1582* (GW1119), at the L4 larvae stage as confirmed by the vulva in the inset. The derepression of the reporter in the gonad is not in germ cells, but does occur in the somatic gonad cells marked with asterisks: DTC (distal tip cells), gonadal sheath, spermathecal cells. **F**, Z-projection of confocal images showing the nuclear GFP derepression of the heterochromatic reporter *pkIs1582* (GW1119) in nearly all if not all somatic cells of an *lsm8^-/-^* worm. Bar, 50 μm. **G**, GFP and DIC merged images at a single focal plan showing the optimal view of germ cells (inside dashed red line), which are not derepressed in *lsm8^-/-^* worm (GW1119) even treated with RNAi against piRNA factors such as *csr-1*.

**Supplementary Figure S3:**
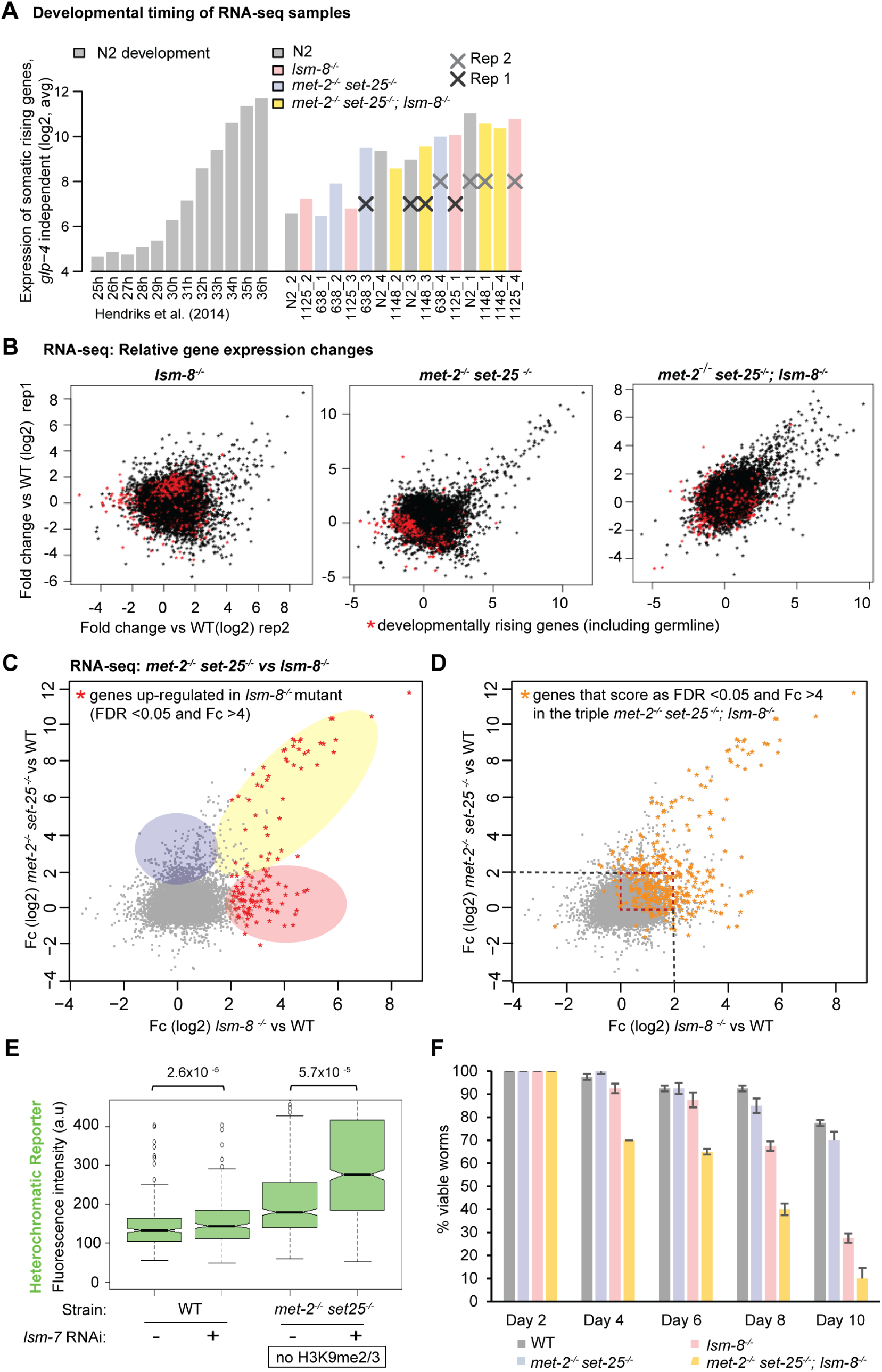
LSM2-8 mediated silencing is independent of H3K9 methylation. **A,** Gene expression data were collected over a time course at 25°C and the average expression of germline genes that were found to increase during this time course is plotted in the left part (Hendricks *et al.,* 2014). This analysis allowed us to compare our samples to the average expression of germline genes and somatic genes that increase naturally within L3 larval stage (see Experimental procedures). Samples from the four different genotypes matched by developmental timing were selected accordingly and assigned to replica 1 and replica 2. Bioinformatics analysis was pursued with these samples. **B,** Relative gene expression profiles as scatter plots. Fold-change (log2) in gene expression of two replicas of RNA-seq from sorted L3 worms of *lsm-8 ^-/-^*, *met-2^-/-^ set-25^-/-^* and the triple (*lsm-8^-/-^*, *met-2^-/-^ set-25^-/-^*) mutant versus WT. Each dot corresponds to a gene. Red dots are rising genes, genes with increased expression level during the time course described (Hendriks et al., 2014), which do not change significantly in any of the mutant strains. **C,** Scatter plot comparing the relative gene expression between the *lsm-8* (x axis) and the *met-2 set-25* double mutant (y axis). Common upregulated genes are shaded yellow; 36% of genes upregulated in the *lsm-8* mutant (FDR <0.05 and Fc >4) are also upregulated (FDR<0.05 and Fc >4) in the *met-2 set-25* mutant. *lsm-8^-/-^*-specific upregulated genes are shaded in pink; *met-2^-/-^ set-25^-/-^-s*pecific are in blue. **D,** Comparison of the *lsm-8* and *met-2 set-25* mutants RNA-seq data, as in (C), overlaid by the set of genes that are upregulated (FDR <0.05 and Fc >4) in the triple mutant *met-2 set-25; lsm-8* (orange stars). The dotted red square highlights genes for which the repression pathways are clearly additive: the orange dots indicate genes that are highly derepressed (>4 Fc) in the triple mutant, but mildly <4 Fc derepressed in either the *lsm-8* or *met-2 set-25* mutant as shown in this graph. **E,** Quantitation of derepression of GFP expressed from the *gwIs4* heterochromatic reporter in L1 progeny in WT and *met-2 set-25* mutant genotypes, respectively from strains GW76 and GW637, after control or *lsm-7* RNAi, displayed as in Fig S1. n= 339, 1004, 426, 673. **F,** Survival assay as in Figure 2. The *met-2^-/-^ set-25^-/-^; lsm-8^-/-^* worms die prematurely compared to the *lsm-8^-/-^* mutant. N=4, n= 40 worms per genotype, bars =s.d.

**Supplementary Figure S4:**
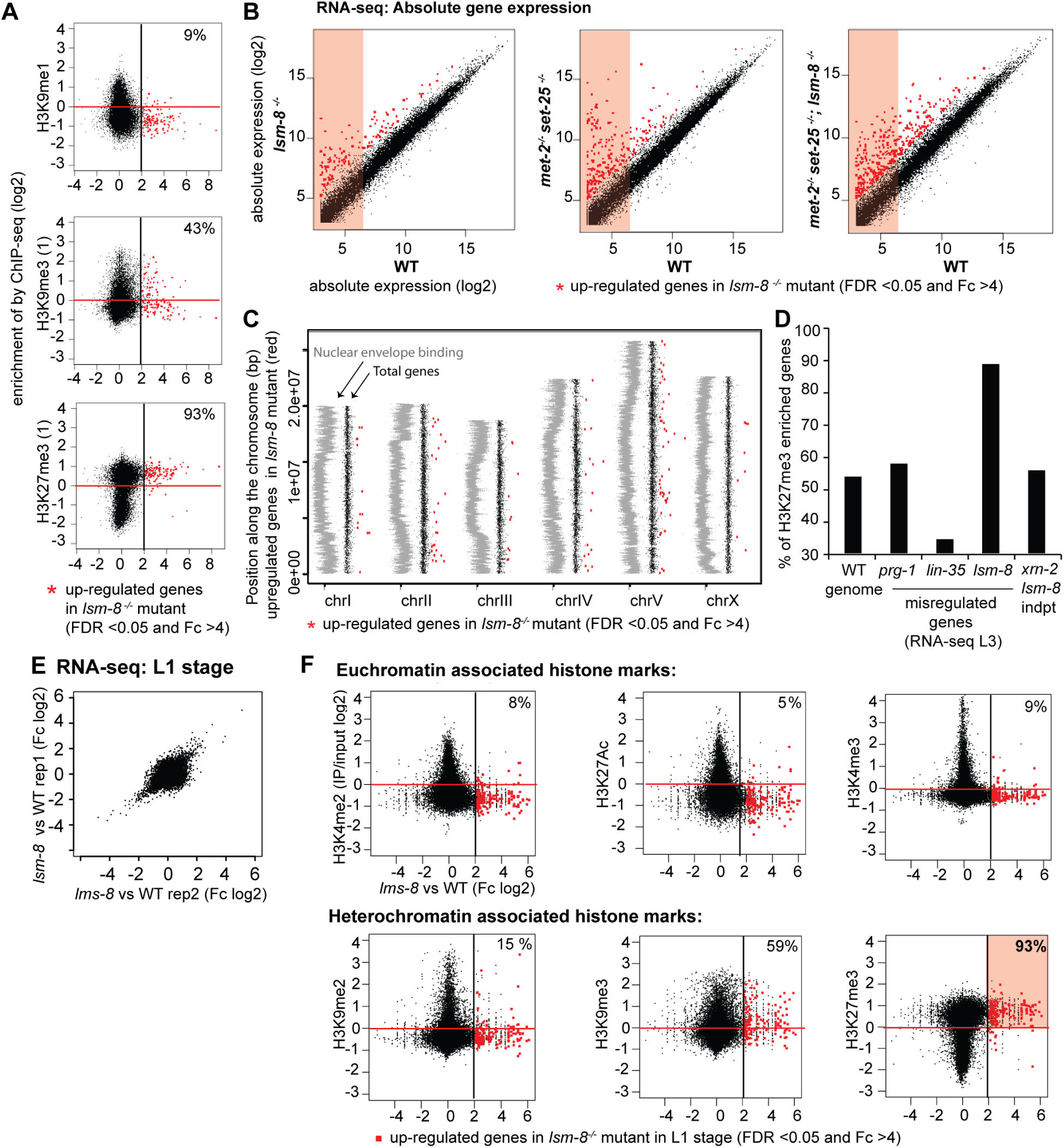
Genes silenced by LSM2-8 carry the Polycomb mark H3K27me3, have a low steady-state expression and are not enriched on chromosome arms. **A,** Correlation of log2(FC) in *lsm-8^-/-^* with the H3K9me1 mark ChIP-seq data and with additional ChIP-seq data for H3K9me3 and H3K27me3 using different antibodies than those used in Figure 3 (see Experimental materials). **B,** Scatter plots comparing absolute transcript abundances (log2 of normalized reads count) of annotated genes in *lsm-8^-/-^*, *met-2^-/-^ set-25^-/-^* and the triple (*lsm-8^-/-^*, *met-2^-/-^ set-25^-/-^*) mutant versus WT. Boxes with pink background indicate low abundance values smaller than 6 in log2 scale for genes considered to be repressed in WT. This corresponds to 64 normalized RNA-seq reads per gene, in contrast to 1024 reads per gene represented by a value of 10. Note the large proportion among the genes upregulated in the assessed mutants (above the diagonal), which are repressed or very poorly expressed in WT. **C,** Distribution of upregulated genes in *lsm-8^-/-^* along chromosomes. LEM-2 ChIP enrichment plotted over chromosomes (embryonic WT data from (Gonzalez-Sandoval et al., 2015)) is in grey, indicating proximity to the nuclear periphery. Up-regulated genes in *lsm-8^-/-^* (FDR <0.05 and Fc >4) represented by the red dots are plotted over autosomes and X chromosome. **D,** Comparison between our RNA-seq and other available RNA-seq datasets (Latorre et al., 2015; Miki et al., 2016; Wang et al., 2014) in L3 stage *C. elegans*, for the percentage of H3K27me3-enriched genes among misregulated genes. We classify a gene as enriched for H3K27me3, if it has positive reproducible enrichment of H3K27me3 over input from two ChIP-seq datasets from ModEncode (Table -S3). Genes upregulated in *xrn-2* RNAi treated worms (Miki et al., 2016) but not upregulated in *lsm-8* mutant worms are not significantly enriched for H3K27me3 (Table S2). **E,** Scatterplots of fold changes of L1 sorted *lsm-8* mutant worms versus WT computed from the normalized read counts per gene in log2 scale for each of two independent RNA-seq replicate sample pairs, as in Fig. 3B. **F,** Scatter plots contrasting on the x axis the gene expression changes in L1 (Fc of sorted *lsm-8* mutant worms versus WT as determined by EdgeR) to the enrichment of the indicated histone mark over input samples (y axis), as in Fig. 3D.

**Supplementary Figure S5:**
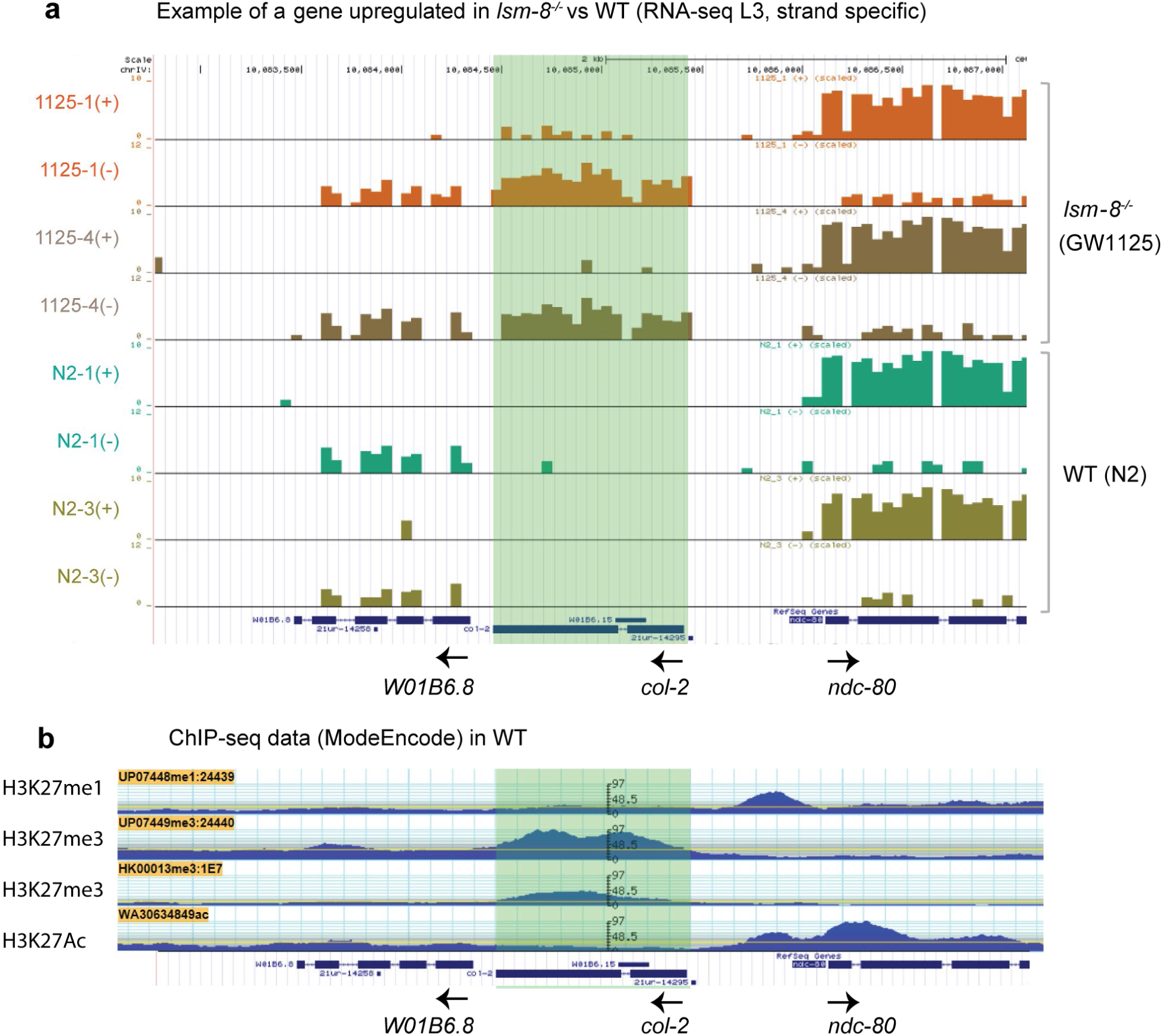
LSM-8 ablation does not alter transcription termination accuracy, strand specificity nor splicing. **A,** UCSC genome browser view showing wiggle tracks from positive (+) or negative (-) strands show the differential expression of the *col-2* gene, which is upregulated in *lsm-8 ^-/-^* compared to WT (y axis in log2). The expression level of the neighboring genes is not affected and termination defects are not observed. All introns were as efficiently spliced in *lsm-8^-/-^* as in WT. **B,** G browse view showing the ModEncode ChIP-seq tracks for H3K27me1, H3K27me3 (two different antibodies) and H3K27Ac at the same genomic locus (IV:10,082,495..10, 087, 496) around the *col-2* gene, as shown in (A). The *col-2* gene is upregulated in *lsm-8 ^-/-^* compared to WT and enriched for H3K27me3, as 95% of the genes upregulated in *lsm-8 ^-/-^*.

**Supplementary Figure S6:**
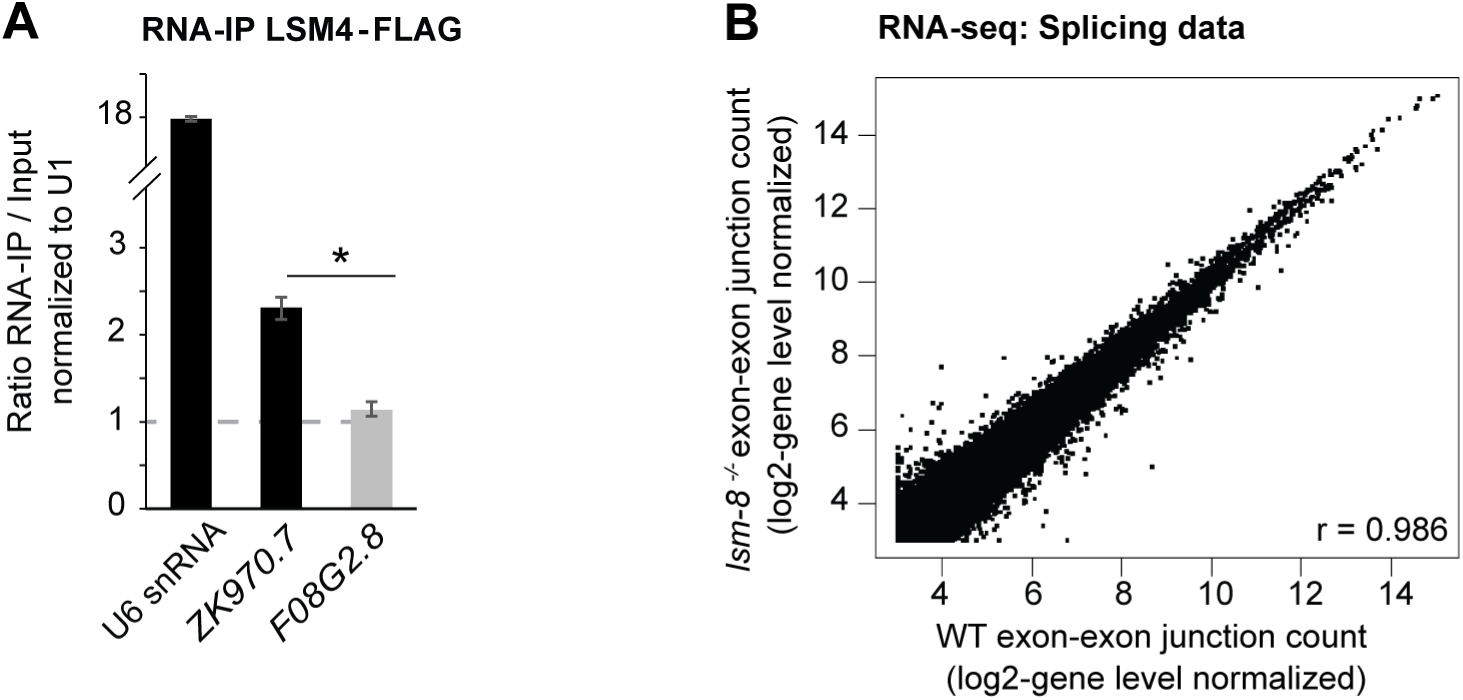
*lsm-8* deletion does not affect splicing globally. **A,** RNA IP-qPCR. LSM-4-FLAG RNA IP analysis in native conditions. RNA levels were normalized to input and U1snRNA levels. *ZK970.7* is upregulated in *lsm-8^-/-^* (*lsm-8* target gene) and associate with LSM4 (>1), whereas *F08G2.8* is not (non-target gene) and do not associate with LSM4. Those two examples suggest that the LSM-8 complex can bind to the RNAs it regulates. N=2, n=3, bars: s.e.m. **B,** Reads which align on exon-exon junctions were counted in *lsm-8^-/-^* and WT worms. Scatter plot compares exon-exon junction mapped reads (log2) normalized to their intrinsic gene level in WT (x-axis) and *lsm-8^-/-^* worms (y-axis).

**Supplementary Figure S7:**
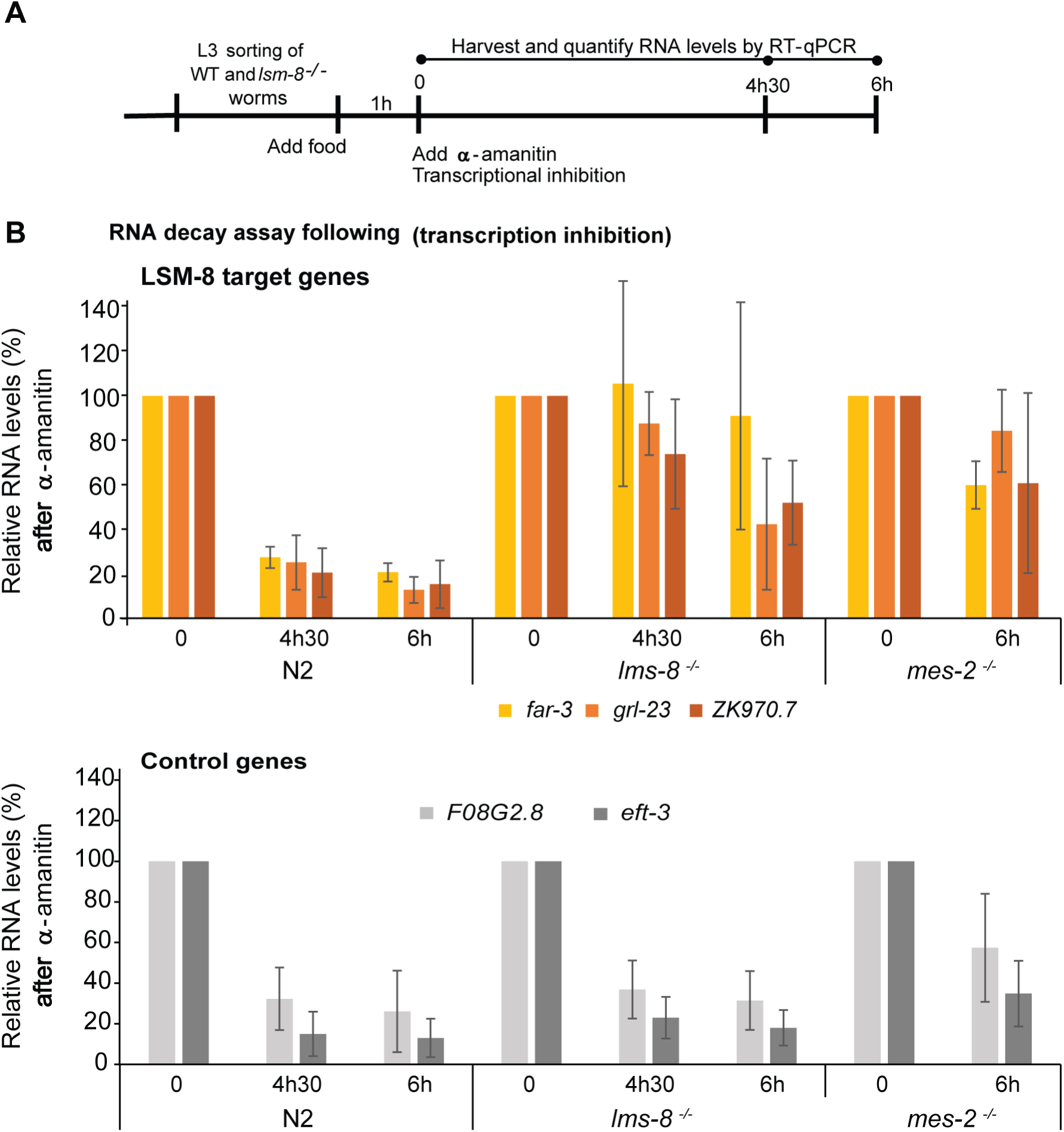
LSM2-8 promotes the degradation of specific transcripts. **A,** Scheme of the RNA decay assay. WT and *lsm-8 ^-/-^* worms were sorted, re-fed with OP50 in liquid culture for 1h at room temperature and treated with 50 μg/ml final concentration of α-amanitin, which inhibits Pol II and Pol III transcription. RNA was isolated at time 0, 4.5h and at 6h, as indicated for each independent experiment. **B,** RNA levels of three transcripts affected by LSM-8 (upper graph) and two control transcripts (expression not affected by LSM-8, lower graph) were determined by RT-qPCR and normalized to 18S rRNA levels which are insensitive to α-amanitin. The value at 0h is defined as 100%. N=3, n=3, bars = s.e.m.

Supplementary Table S2: RNA-seq data for genes in *lsm-8*, *met-2 set-25* and *lsm-8, met-2 set-25* triple mutant versus WT at the L3 stage. (Excel file uploaded separately)

Supplementary Table S3: Enrichment of histone marks in L3 (ModEncode Data, Excel file uploaded separately)

Supplementary Table S4: Lists of biological processes affected in the different mutants (GO analysis). (Excel file uploaded separately)

Supplementary Table S5: RNA-seq data for genes in *lsm-8* mutant versus WT at the L1 stage. (Excel file uploaded separately)

## Experimental Procedures

### Worm strains and growing conditions

Table S1 lists the strains used in this study. Strains with deletion alleles and reporters obtained from the *C. elegans* knockout consortium or made by the CRISPR/Cas9 system were outcrossed 2 to 6 times to the N2 (WT) strain. Worms were grown on OP50 and maintained at 22.5°C, except when frozen or manipulated at room temperature (RT).

The *lsm-8* deletion allele xe17 (sequence below) was generated by replacing the entire coding sequence of the *lsm-8* gene with the red pharynx marker [*myo2p::mCherry::unc54 3’UTR*] using an adapted version of the CRISPR/Cas9 technique (Katic et al., 2015). For this, the N2 worms were injected with the following mix pDD162 (Cas9, Dickinson et al., 2013) 100 ng/μl, LSM8 sgRNA1 (Fwd) in PIK111 100 ng/μl, LSM8 sgRNA3 (Rev) in PIK111 100 ng/μl, the indel plasmid *lsm-8Δ*-mCherry in pIK37 100 ng/μl and *Pmyo-3::gfp* 5 ng/μl.

### DAPI staining and live microscopy

DAPI staining was carried out on WT and *lsm-8^-/-^* (handpicked) worms from different developmental stage (not mixed) and mounted on poly-L-lysine coated slides. Two independent biological replicates were performed. The freeze cracking of worms by liquid nitrogen in Eppendorf tubes was followed by fixation for 5 min in methanol at -20°C, and 2 min in 1% paraformaldehyde at rt for all stages. After fixation, 3 x 5 min washes with PBS supplemented with 0.25% TritonX100 (PBSX) were done with the last wash optionally lasting overnight (ON) at 4°C. DAPI (1μg/ml) was added for 10 min at rt and was washed twice before mounting the slides with n-propyl gallate. For live imaging, animals were mounted on slides coated with 2% agarose pads, supplemented with 0.1% sodium azide and 1mM levamisole, in most cases.

Microscopy was carried out on a spinning disc confocal microscope (AxioImager M1 [Carl Zeiss] + Yokogawa CSU-22 scan head, Plan-Neofluar 100×/1.45 NA oil objective, EM-CCD camera [Cascade II; Photometrics], and VisiView 2.1.4 software, (Fig. S1D,E, Fig. 2C, Fig. 5B, Fig. S2B-E) either Axo imager 2.1 Zeiss, (Fig. 2D, Figs. 2I-Ll, 3A-D). Images, 3D reconstruction (maximum intensity Z-projections) and fluorescence intensity analysis were generated using Fiji/ImageJ software.

### RNAi experiments

RNAi was performed at 22.5°C by placing synchronized L1 worms on feeding plates as previously described (Timmons et al., 2001). Synchronized L1 larvae were obtained by bleaching gravid adults and the eggs recovered were left to hatch overnight at RT in M9. All RNAi clones used against LSM complexes subunits and used in the targeted RNAi screen were sequenced and a blast analysis performed first to confirm the specificity of the targets. At least, three independent biological replicates were performed for each RNAi experiment. As a mock RNAi control, the L4440 vector (Fire vector library) was modified by removing an *EcoRV* fragment containing 25b.

For RNAi against *xrn-2*, bacteria expressing dsRNA were diluted with mock RNAi bacteria to feed the GW306 and GW1119 strain in order to get a milder phenotype and thus enough progeny in which to assess derepression. Both *lsm-8* heterozygous and homozygous worms (GW1119) were subjected to RNAi treatment, but only homozygous worms were used to assess the RNAi effect. For the RNAi with LSM-8 potential co-factors, most of the chosen candidates were LSM2-8 subunits related or controls. Co-regulated genes were predicted though a clustering analysis in SPELL (http://spell.caltech.edu:3000/). List of interacting partners were predicted through the Wormbase. The derepression was assessed by the worm sorter as described in Figure 1 for RNAi hits that produce L1 larvae in the next generation. For RNAi hits that caused larval arrest or embryonic lethality, derepression was assessed by microscopy using in each case, adequate controls.

### Quantitation of derepression

Derepression was scored at specific developmental stages by fluorescence microscopy using standardized exposure and illumination conditions. Quantitation of GFP intensity in different conditions was done using Fiji/ImageJ software and the ROI manager, for semi-automated analyses. The fluorescence intensities from whole animals at similar developmental stages were also compared.

Quantitation of derepression by the worm sorter, COPAS BIOSORT (Union Biometrica), was performed in L1 worms according to manufacturer’s guidelines. Visual inspection of the selected and monitored worms showed that >99% of all worms matched the size criteria. Data corresponding to the fluorescence intensity (PH Green or PH Red) were analyzed and plotted in boxplots using R studio. The EXT (1-5) was extracted to exclude possible remaining bacteria.

### Survival assay

Worms of indicated genotypes were synchronized by bleaching, and when they reached the L4 stage (Day 2 at 22.5°C), ten worms were isolated onto plates containing OP50 bacteria. Four independent biological replicates were performed. The number of worms alive was determined every 24h. At Day 4, surviving adults worms from each genotype (even sterile ones, *lsm-8^-/-^* and *met-2 set-25; lsm-8^-/-^*) were transferred to a new plate to avoid contamination with the progeny and at Day 6, only adults of WT and *met-2 set-25* strains were transferred, since the other sterile worms were too fragile to move without being killed.

### Chromatin Immunoprecipitation (ChIP) experiments

∼20,000 WT and *lsm-8*^-/-^ homozygous L3-L4 larvae stage were isolated using the COPAS BIOSORT instrument (Union Biometrica), according to manufacturer’s guidelines. Three independent biological replicates were performed. Visual inspection of the sorted worms showed that >90% of all worms were expressing appropriate markers (*i.e.*, red fluorescence but no GFP expression in the pharynx for *lsm-8^-/-^*, and no markers for the WT) and 90% matched the desired size and morphological criteria that corresponds to the stage of interest.

Antibodies used for the ChIP were rabbit anti-H3K27me3 (ChIP, Millipore, 07-449), whose specificity was confirmed by peptide binding, and IF on a *mes-2* mutant (data not shown).

H3K27me3 ChIP was performed as previously described (Zeller et al., 2016). In brief, chromatin was incubated overnight with 3 µg of antibody coupled to Dynabeads Sheep Anti-Rabbit IgG (Invitrogen), in FA-buffer (50 mM HEPES/KOH pH7.5, 1 mM EDTA, 1% Triton X-100, 0.1% sodium deoxycholate, 150 mM NaCl) containing 1% SDS. Chromatin/ antibody complexes were washed with the following buffers: 3 x 5 min FA buffer; 5 min FA buffer with 1M NaCl; 10 min FA buffer with 500 mM NaCl; 5 min with TEL buffer (0.25 M LiCl, 1% NP-40, 1% sodium deoxycholate, 1 mM EDTA, 10 mM Tris-HCl, pH 8.0) and twice for 5 min with TE. Complexes were eluted at 65°C in 100 µl of elution buffer (1% SDS in TE with 250 mM NaCl) for 15 min. Both input and IP samples were incubated with 20 µg of RNAse A for 30 minutes at 37°C and 20 µg of proteinase K for 1 h at 55°C. Crosslinks were reversed overnight at 65°C. DNA was purified using a Zymo DNA purification column (Zymo Research).

### RNA-IP in native conditions

Enriched L3 stage worms (GW1004 which contains extrachromosomal arrays expressing LSM-4-GFP/3xFLAG-tagged from a fosmid which was obtained from the “*C. elegans* TransgeneOme” consortium) were collected as 300-500 μl of pelleted worms and lysed at 4°C with a Dounce Tissue Grinder (150 strokes for each 500 μl, BC Scientific, Miami, FL, USA) in an equal volume of lysis buffer (30 mM HEPES/KOH pH 7.4, 100 mM KCl, 1.5 mM MgCl_2_, 0.1% Triton X-100, Protease Inhibitor Cocktail Tablets, EDTA-free, Roche Rnase inhibitor, rRNAsin 1.25μl/ml of lysis buffer). Lysates were cleared at 16 000 x *g* for 15 min. 4 mg of lysate proteins were incubated with 40 μl of pre-washed anti-FLAG M2 magnetic beads (Sigma– Aldrich) for 2 h. Washes were performed in lysis buffer. For RNA extraction, washed magnetic beads were resuspended with 100 μl of lysis buffer and 400 μl Trizol® (Ambion) and the samples were snap-frozen in liquid nitrogen. Two independent biological replicates were performed.

### RNA extraction

For the RNA-seq experiment WT, *met-2 set-25*, *lsm-8*^-/-^, and *met-2 set-25; lsm-8^-/-^* worms were isolated using the COPAS BIOSORT instrument according to the fluorescent criteria (non-green pharynx worms) using the size criteria of L3 stage larvae in 4 independent biological replicates. For L1 RNA-seq experiment, worms were synchronized prior to the sorting process.

Synchronized L1 larvae were obtained by bleaching gravid adults and the eggs recovered were left to hatch 16h at RT in M9.The isolation of WT and *lsm-8*^-/-^ L1 larvae was made similarly with the fluorescent criteria (non-green pharynx worms) and the size criteria of L1 stage larvae. The larvae were refeed for 2.5h after the sorting process. For all RNA based experiments, before RNA extraction, worms were washed 3x in M9 and re-suspended in 100μl of M9, 400μl of Trizol® (Ambion) and snap-frozen in liquid nitrogen.

Extraction of RNA used 4 freeze-thaw cycles from liquid nitrogen to a 42°C heat bath, followed by the addition of 200μl of Trizol® to each sample. Vigorous vortexing at room temperature (rt) in 5 cycles (30 sec vortex, 30 sec on ice), was followed by 5 min at rt. RNA extraction was with 140μl chloroform, vigorous shaking for 15 sec, and 2 min at rt. The samples were centrifuged at 12000 rcf at 4°C, and the aqueous phases were transferred to fresh tubes. An equal volume of 70% EtOH was added slowly and the homogeneous mixture was transferred to a Qiagen RNeasy spin column (RNeasy kit, QIAGEN 74104). QIAGEN protocols including a subsequent 30 min DNAse treatment. For L1 RNA-seq samples, the extraction was done using the Zymo DirectZol microRNA kit (R2060).

### RT-qPCR

Primers were designed to be exon-junction spanning where possible, and are listed below. cDNA synthesis was performed using the (AMV cDNA kit, NEB, E6550S) according to the manufacturer’s protocol using random primers and 0.1-3 µg of total RNA per sample according to the experiment. qPCR was performed on a StepOnePlus real time PCR system (Applied Biosystems) using SYBR Green Mastermix (Applied Biosystems; 4309155). Further analysis was done in Microsoft Excel. All primer pairs were tested and selected for amplification efficiencies ranging from 85-100%. For gene expression analysis in Fig. 1 and S1, ΔΔCT method was used, *his-56* and *pmp-3* were used for sample normalization. For ChIP-qPCR, sample data were normalized to corresponding input chromatin. Candidate genes were chosen in Fig. 6 based on their expression changes and on their enrichment for H3K27me3 in WT worms. For RIP-qPCR in Fig. S6A, RNA levels were normalized to corresponding input and to the U1snRNA levels.

### RNA decay assay

WT, *lsm-8^-/-^* and *mes-2^-/ -^* (F2) L3 larvae were sorted and re-fed with OP50 in liquid culture for 1 h at RT. Subsequently α-amanitin (Sigma-Aldrich) was added to a final concentration of 50 mg/ml, to block transcription and stall larval development (Miki et al., 2014). About 750 worms were harvested in duplicate in each of the three independent biological replicates, and for each sampling point. They were washed twice with M9 medium, resuspended in 400 ml of Trizol® (Life Technologies) and frozen in liquid nitrogen. To assess the RNA decay, RNA levels of genes affected or not by the LSM2-8 complex (expression level) were quantified before and after the transcriptional inhibition in each genotype. LSM-8 target genes were selected by their higher expression levels in *lsm-8^-/-^* versus WT (RNA-seq), and their enrichment for H3K27me3 in L3 larvae, yet it was desired to have detectable levels in WT control. In this assay, cDNA was generated from total RNA by the SuperScript III First-Strand Synthesis System (Thermo Fisher Scientific) using random primers and the 5x FS buffer for better yields. Three micrograms of total RNA were used as a template for reverse transcription reaction (20μl), and 0.66μl of the reaction was used for qPCR reaction (10μl). RT-qPCR for this assay was performed using PowerUp SYBR Green Master Mix (Thermo Fisher Scientific), specific primers for mature/spliced mRNAs (complementary to an exon-exon junction; *grl-23, F08G2.8*) or for pre- and mature mRNAs (*far-3, ZK970.7*) or for pre-mRNA only (*eft-3*) and using StepOnePlus Real-time PCR Systems (Applied Biosystems) according to the suppliers’ protocols. For primer sequences for *eft-3* and 18S ribosomal RNA, see (Miki et al., 2014). Because pre-mRNA levels are expected to be more directly affected by transcription inhibition, *eft-3* pre-mRNA was used by us and by others (Miki et al., 2014) as a control for the efficiency of the α-amanitin treatment in inhibiting transcription. The high expression levels of *eft-3* makes it an adequate control to verify the potential extent of the transcriptional inhibition. In addition, *eft-3* is also a control gene in the sense that it is not regulated by *lsm-8*.

### RNA-seq

Total RNA was treated for the L3 samples additionally with the Turbo DNA free kit (Ambion, AM1907), depleted for rRNA using Ribo-Zero Gold kit from Epicentre and depletion validated through Agilent Bioanalyzer analysis. Subsequent library preparation was performed with a ScriptSeq v2 RNA-Seq library preparation kit, stranded (Epicentre). Library preparation for the L1 samples was performed with the TrueSeq Total RNA preparation kit, stranded (Illumina).

The quality of the resulting libraries was assessed with an Agilent Bioanalyzer and concentrations were measured with a Qubit fluorometer prior to pooling. 50 bp single-end sequencing was done on an Illumina HiSeq 2500.

### Processing of the RNA-seq and ChIP-Seq data

The RNA-seq samples from four independent biological replicate samples L3 were mapped to the *C. elegans* genome (ce6) with the R package QuasR v1.22.0, (www.bioconductor.org/packages/2.12/bioc/html/QuasR.html) with the included aligner bowtie (Langmead et al., 2009) considering only uniquely mapping reads for mRNA. The command “proj <-qAlign(“samples.txt”,“BSgenome. Celegans.UCSC.ce6”)” instructs bowtie to align using the parameters “-m 1 --best --strata --phred33-quals”. Since the used replicas differed slightly in timing (Fig. S3A), we incorporated a blocking factor in the linear model treating the replicates as different batches. For splice junction quantification we used the spliced alignment algorithm SpliceMap (Au et al., 2010). The command used was “proj <-qAlign(“samples.txt”,“BSgenome.Celegans. UCSC.ce6”,splicedAlignment=TRUE)”. The command to create various count tables was qCount(proj,exons,orientation=“same”). For gene quantification, gene annotation from WormBase was used (WS190). The EdgeR package v 3.24.0 was used to determine fold changes (Fc) and false discovery rates (FDR) of differential transcript abundances. The repeat element quantitation was based on UCSC (genome.uscsc.edu) repeat annotation. To normalize for sequencing depth, each sample was divided by the total number of reads and multiplied by the average library size. Transformation into log2 space was performed after the addition of a pseudocount of 8 in order to minimize large changes in expression caused by low count numbers. The various count tables used throughout this study were normalized separately. To determine the developmental timing of each RNA-seq sample, we previously used a set of 2050 genes shown to gradually rise between 25h and 36h post hatching at 25°C (all rising genes)(Hendriks et al., 2014). While most of those genes are germline genes and thus stop being expressed in *glp-4* mutants which are devoid of germ cells (Hendricks *et al.,* 2014, Fig. S3A), we noticed that a subset of those rising genes (n=162) actually still continued to rise even in *glp-4* mutant worms (Hendricks *et al.,* 2014, Fig S3A). We therefore split the 2050 genes into two separate groups, a germline developmental signature (n=1888) and a somatic developmental signature (n=162) and used the latter to infer developmental timing (Fig S3A). We got the same result using all rising genes (data not shown). To quantify potential changes in splicing in *lsm-8^-/-^* as opposed to WT, we quantified the expression of all the exon-exon junctions from the spliced alignments using no annotation. The command used to create the exon-exon junction count table was qCount(proj2,NULL,reportLevel=“junction”). These junction counts were then normalized for library size (as described above) and overlapped with gene annotation to assign them to their host gene. Junctions overlapping multiple genes were discarded. The assignment to the host gene was then used to correct the junction expression levels for differences in gene expression. This was done by dividing the junction counts of either WT or *lsm-8^-/-^* by the respective gene expression change depending on the direction of the change. This procedure ensured that junction counts were always deflated and not inflated by the gene expression correction. Finally a pseudocount of 8 was added and the data were log2 transformed. We specifically chose to not use reads overlapping intronic sequences for this analysis as they can reflect changes in mRNA transcription (Habacher et al., 2016) and thus would potentially complicate the interpretation of those results in the light of alternative splicing. The RNA-seq L1 samples were mapped to the *C. elegans* genome (ce10) and processed otherwise as mention above (no blocking factor applied, as for L3). The ChIP-seq data for L3_H3K9me1/2/3 (5036, 5050, 5037, 5040), L3_H3K27me3 (5045, 5051), L3_H3K27ac (5054), L3_H3K4me2/3 (5055, 3576) were downloaded from ModEncode (http://data.modencode.org/) and mapped to ce6 and ce10 using bowtie considering only uniquely mapping reads. Quantitation for each gene was performed by counting the reads overlapping the gene-body. All samples were normalized for total library size, log2 transformed after adding a pseudocount of 8 and and Fc enrichments (log2) were calculated by subtracting the log2 transformed values of the specified input sample (3576, Rep-1) from each ChIP-seq sample.

Misregulated genes in the *prg-1* and *lin-35* mutants (Latorre et al., 2015; Wang et al., 2014) were converted into WB gene names through the Gene ID conversion tool (DAVID), and the resulting genes were compared to their enrichment in H3K27me3 similarly as for the misregulated genes in the *lsm-8* mutant (Table S3).

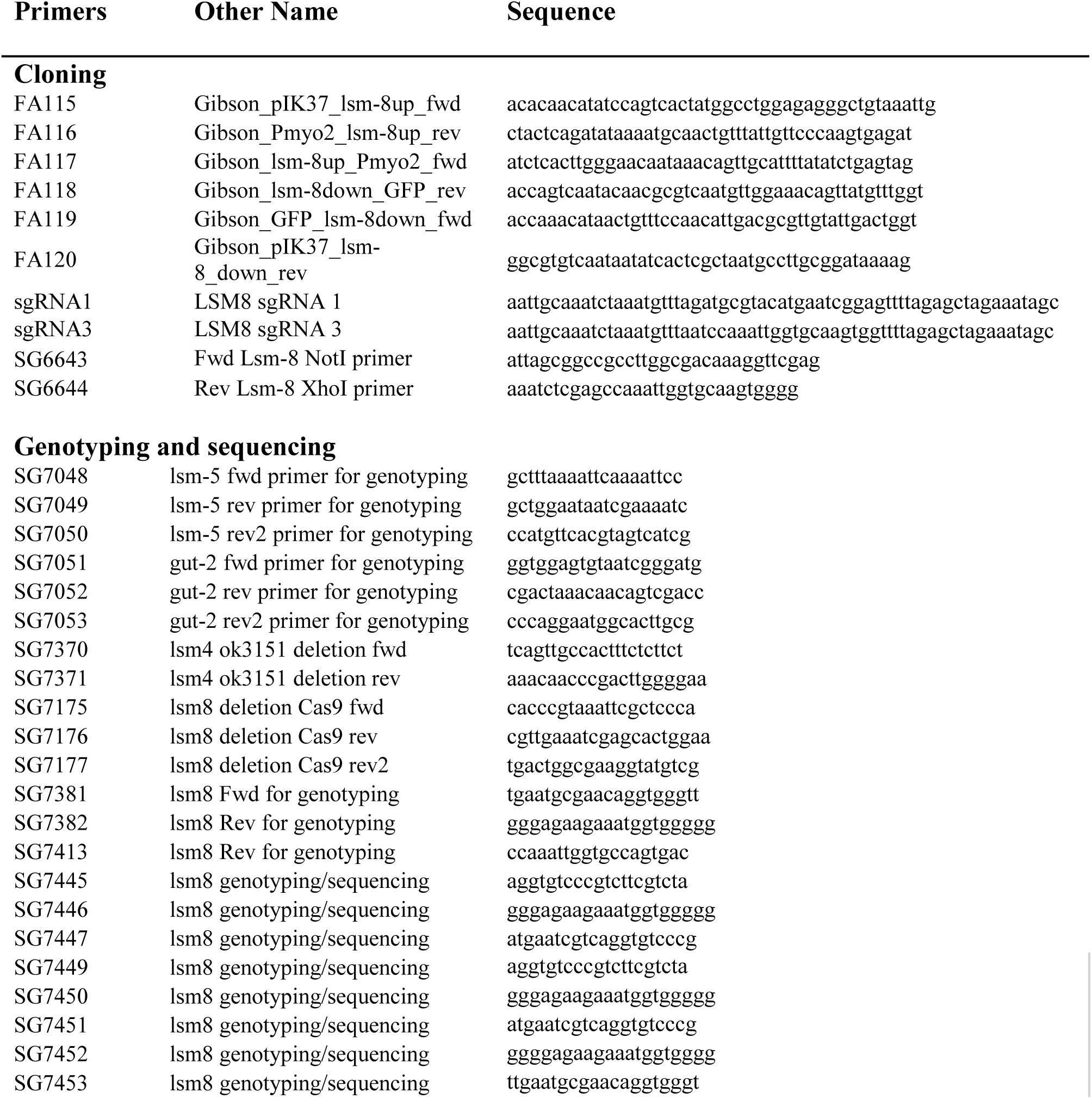
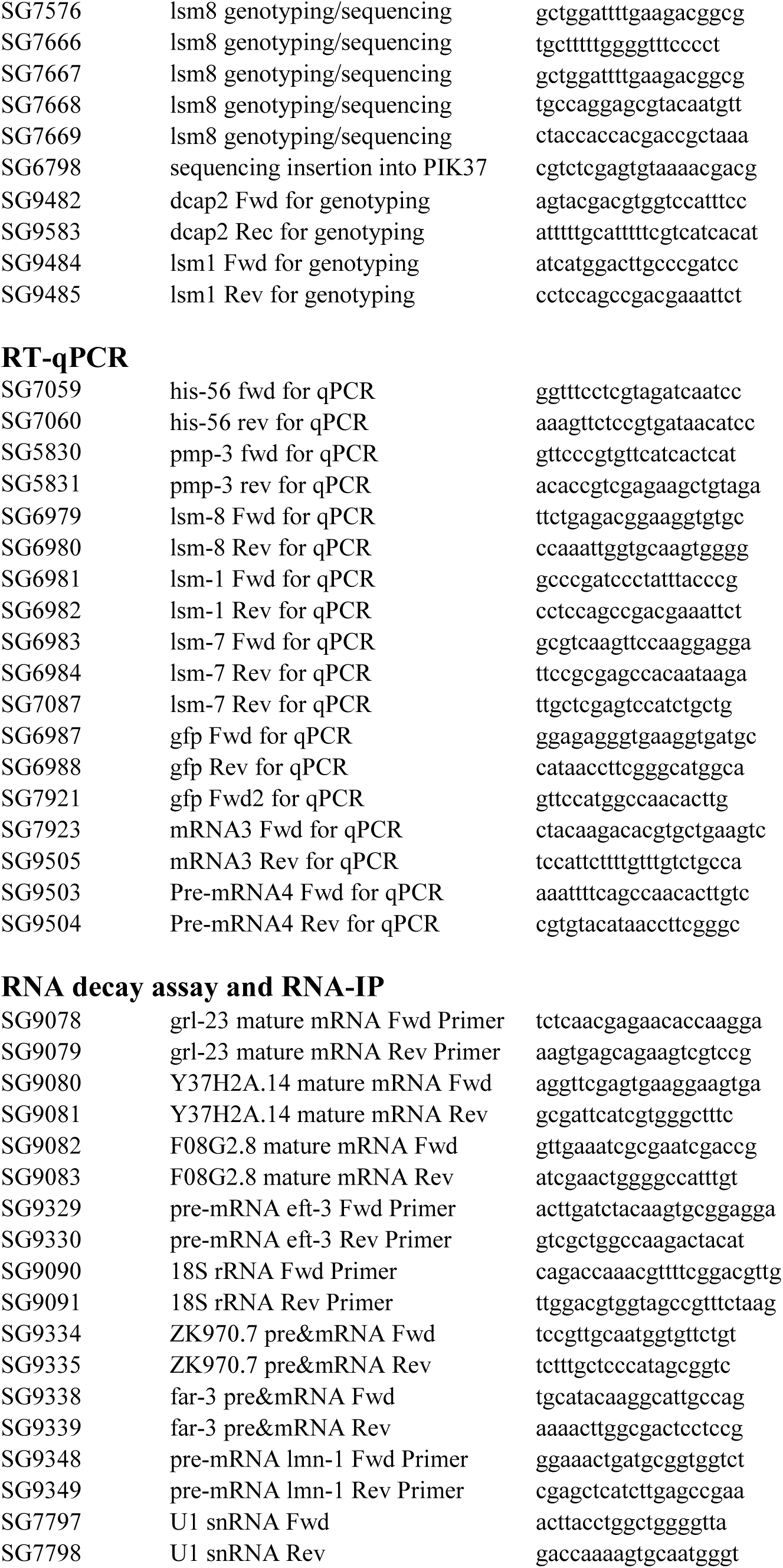
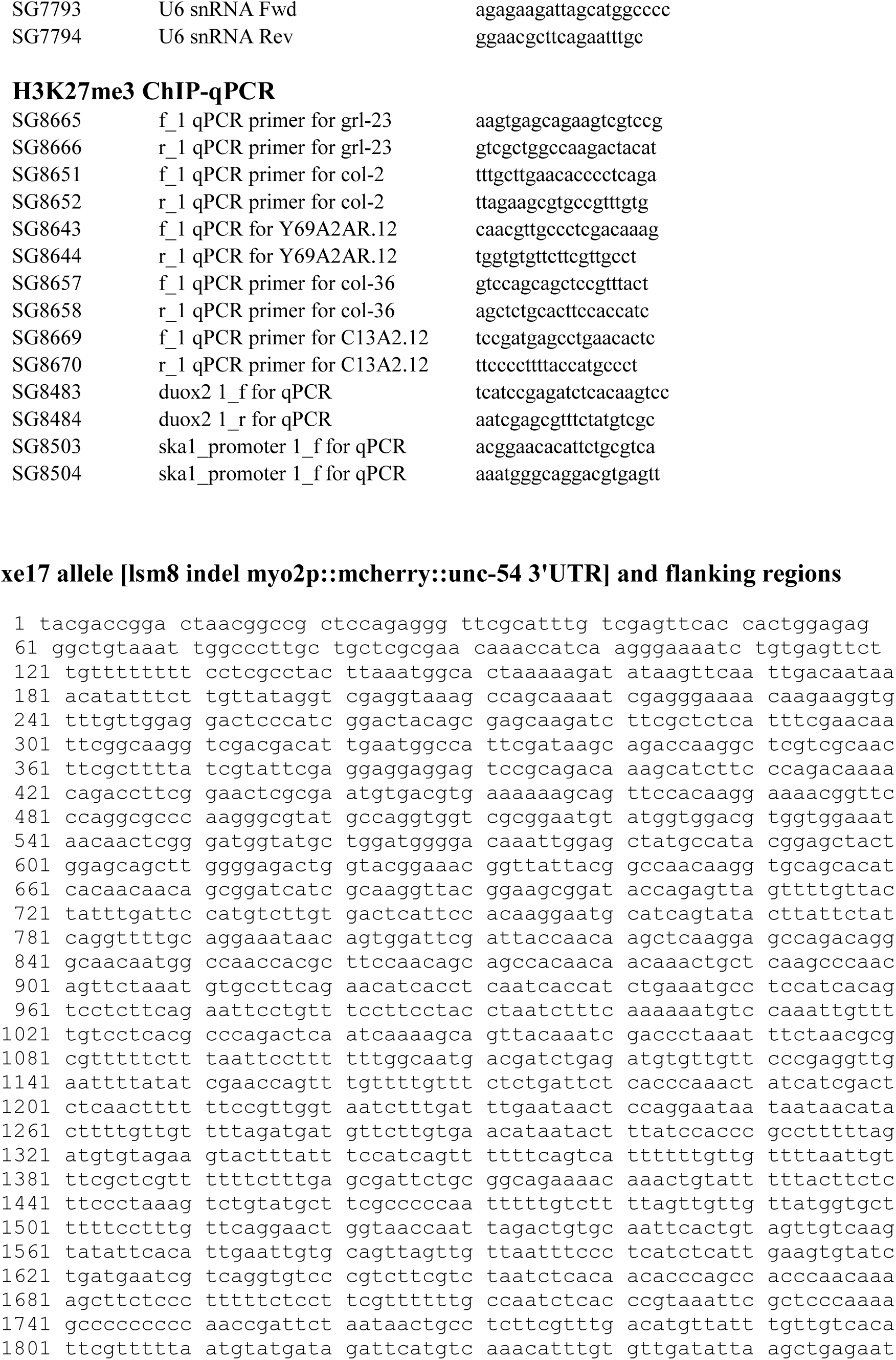
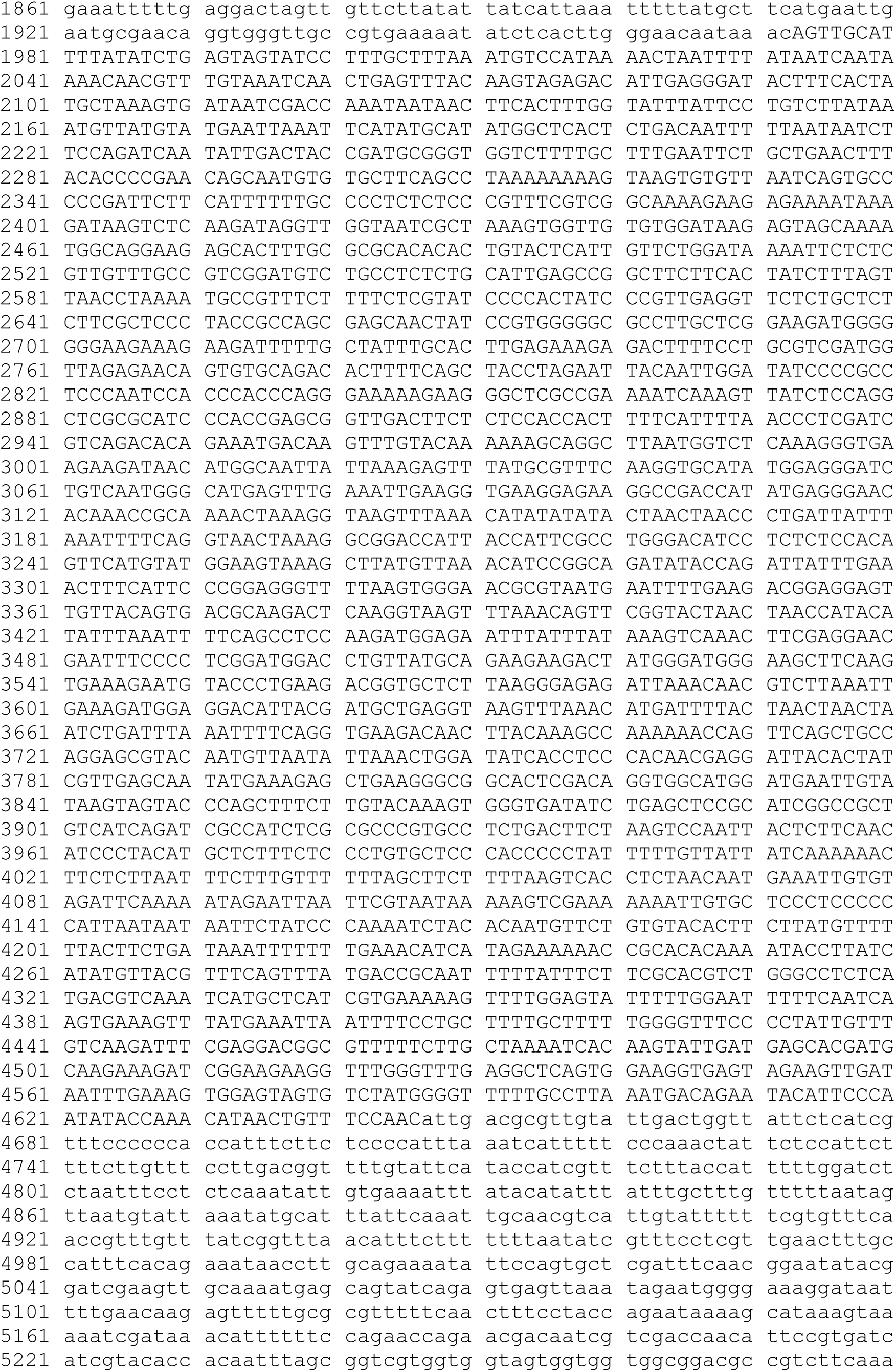
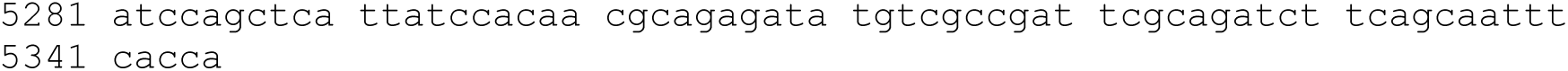
Primers used in this study

